# Proteogenomics of *Blumeria hordei* supports RNA and protein coding innovative potential derived from transposable elements

**DOI:** 10.64898/2026.05.04.722472

**Authors:** Xinyi Liu, Luzie U. Wingen, Alexandros G. Sotiropoulos, Sadegh Balotf, Levente Kiss, Benedikt Schiestl, Veronika Schmitt, Daniela Scheikl, Sonja Dunemann, Nafiseh Sargheini, Bruno Hüttel, Amirhossein Sakhteman, Miriam Abele, Christina Ludwig, Aurélien Tellier, Marion C. Müller, Ralph Hückelhoven

## Abstract

Some filamentous plant-pathogenic fungi have comparably large genome sizes within the fungal kingdom due to the proliferation of transposable elements (TEs). *Blumeria hordei* (*Bh*), the causal agent of the powdery mildew disease on barley, is a filamentous obligate biotrophic fungus. Compared to other ascomycetes, it contains a low number of genes but a high genomic TE content of approximately 75%. Yet, a comprehensive understanding of the contribution of TEs to the RNA and protein landscape of *Bh* is lacking. Here, we use *Bh* as a model to study transcripts and proteins derived from genes and individual TEs. Therefore, we created two high-quality genome assemblies of the German *Bh* isolate TUM1 and the Australian *Bh* isolate AUS1. We applied deep proteomics with mass spectrometry, long-read and short-read sequencing on both DNA and RNA. Based on these multi-omic resources, we completed nearly gapless genome assemblies, new gene and TE annotations, and effector predictions. Using long-read RNA sequencing, we detected extensive co-transcription of TEs and genes as TE-gene chimeric transcripts. We identified previously unpredicted splice variants or genes, partially supported by proteomics. The intergenic and TE genomic space of *Bh* TUM1 gives rise to thousands of transcripts and several novel TE-derived proteins that lack from previous TE protein predictions. Together, this supports an existing potential for expression of novel transcripts and proteins from highly abundant TEs in the *Bh* genome.

## Introduction

Fungal plant pathogens have very diverse sizes and structures of genomes. However, they all encode for effector proteins that play crucial roles in enabling and promoting successful infections. Upon penetration of the plant cell wall, parasitic fungi deliver effectors from invasive hyphal structures or specialized haustoria into the plant apoplast or cytoplasm. Such effectors often suppress the plant’s immune system by interacting with host targets that function in pathogen recognition or defense. Many such targets act in basal disease resistance that largely builds on pattern-triggered immunity (PTI), in which the plant recognizes and responds to molecular patterns of the pathogen or is released upon pathogen activity. In addition, effectors target plant susceptibility factors that support pathogen demands in accommodating infection structures or metabolic requirements. In contrast, plant host genomes carry resistance genes (*R* genes), which allow the recognition of pathogen strain-specific effectors (avirulence effectors, AVR) and the activation of strong defense mechanisms such as the hypersensitive response to control disease development (Panstruga and Dodds, 2009; Martel et al., 2021; Jones et al., 2024). The mutual influence on each other of ETS and ETI is supposed to drive the rapid evolution of effector genes as they need to adapt to overcome resistance conferred by cognate R genes, leading to frequent gain and loss and a high diversity of effectors in plant pathogens (Franceschetti et al., 2017; Fouché et al., 2018; Martel et al., 2021; Müller et al., 2026).

*Blumeria hordei* (*Bh*), an obligate biotrophic ascomycete, is the causal agent of barley powdery mildew, an important cause of barley yield loss and change in grain quality. *Bh* has both asexual and sexual life cycles, in which the asexual life cycle forms conidia spores that are dispersed by wind, and this is the main form of disease spread in the field (Limpert et al., 1999; ZHANG et al., 2005). Some barley and wheat powdery mildew AVRs are one-to-one recognized via multi-allelic *R* genes, exemplified by powdery mildew genes (*Pm*) in wheat and mildew locus a (*MLA*) in barley(Alam, 2011; Bourras et al., 2015; Lu et al., 2016; Saur et al., 2019; Bauer et al., 2021). Avirulent Blumeria races have the potential to escape the recognition from *R* genes and regain virulence by AVR gene diversification or loss. This and adaptation to diverse host targets may have caused enhanced frequencies of gene duplications and losses in *Blumeria* effector families and higher turnover rates compared to other functional genes (Menardo et al., 2017; Kusch et al., 2024). The identification of the effector repertoire in *Bh* very much advanced through *in silico* prediction tools and genome sequencing (Spanu et al., 2010; Wicker et al., 2013; Frantzeskakis et al., 2018; Müller et al., 2019; Sperschneider and Dodds, 2022; Bilstein-Schloemer et al., 2025). From the sequence analysis, three commonly occurring amino acids (tyrosine, tryptophan, and cysteine) at the N-terminus of many *Bh* effectors were identified, known as the Y/F/WxC motif (Godfrey et al., 2010). Around 500 Candidate Secreted Effector Proteins (CSEPs) were predicted to be encoded by the genome of *Bh* based on Markov clusters (Pedersen et al., 2012; Sabelleck et al., 2025). Many sequence-diverse effector genes encode proteins with RNase-like folding, including a large proportion of CSEPs and *AVR* effectors; some of these encode RNase-Like Proteins Associated with Haustoria (RALPHs) (Spanu, 2017). Well understood effectors that structurally link to RALPHs include *AVR_PM2_* and *AVR_A6,A7,A10/A22,A13_*, which could be a consequence of neo-functionalization of ancient RNase (Praz et al., 2017; Cao et al., 2023; Lawson et al., 2025).

Effectorome annotation, however, extends beyond the characterized effector structural features, as effectors have diverse evolutionary origins, are poorly conserved across families, and might not always contain a signal peptide or common structural scaffold (Bourras et al., 2018). Indeed, two recent studies have identified two effectors that lack a predicted signal peptide and belong to a larger family of comparably big effector proteins composed of two major domains of unknown function (Weiß et al., 2025; Kunz et al., 2026)) Some effector candidates in *Bh* were found to arise from transposable elements (TEs), mobile genomic elements with high copy numbers that are ubiquitously present in eukaryotic genomes. Transposons that translocate by RNA transcripts are defined as class I transposons (retrotransposons) while transposons that translocates as DNA are defined as class II transposons (DNA elements) (Wicker et al., 2007). The TE orders detected in *Bh* are long terminal repeats (LTRs, class I), long-interspersed nuclear elements (LINEs, class I), short-interspersed nuclear elements (SINEs, class I), and inverted terminal repeats (TIRs, class II) (Qian et al., 2023). Both LINEs and LTRs encode proteins such as reverse transcriptase, whereas SINEs do not encode canonical TE proteins and might have diverse origins related to transfer RNA, 7SL RNA, and 5S ribosomal RNA (Kramerov and Vassetzky, 2011). An effector family homologous to the postulated *AVR* genes *AVR_K1_* and *AVR_A10_* (EKA) phylogenetically clusters with first open reading frame (ORFs) of LINEs (*Satine* and *Kryze* LINEs) (Amselem et al., 2015). The *Bh* effector candidate ROPIP1 is encoded on the first ORF of a transposon, which belongs to *Eg_R1* family of SINEs. ROPIP1 interacts with the barley ROP GTPase and susceptibility factor *Hv* RACB and leads to the disorganization of the host microtubule cytoskeleton, which may explain its function in pathogen virulence (Nottensteiner et al., 2018). Although the originally postulated avirulence gene function of some EKAs appears outdated by identification of an *MLA10*-matching avirulence effector, EKAs appear to exert virulence function (Saur et al., 2019). Indeed, targeting them by host-induced gene silencing limits fungal infection success and can be complemented with the expression of RNAi-insensitive protein-coding wobble-constructs (Nowara et al., 2010).

TEs can also have other functions in creating genetic diversity and thus likely contribute to virulence and adaptation. They disrupt genes, drive structural variation, and be co-opted by hosts for neofunctionalization, including exonization, which yields novel, alternatively spliced isoforms and expands protein diversity across eukaryotes (Sela et al., 2010; Alzohairy et al., 2013; Oliveira et al., 2023; Arribas et al., 2024). TEs are closely related to the virulence of plant pathogens. Although TEs can disrupt essential genes, can turn genes deleterious and cause genome instability, they also create diversity to maintain virulence, or to generate new phenotypes (Möller and Stukenbrock, 2017; Fouché et al., 2022). TEs comprise great parts of the genome in pathogens and enhance genome plasticity by driving structural rearrangements in *Zymoseptoria tritici* and enabling horizontal transfer of effector genes such as *ToxA* (Hartmann et al., 2017; McDonald Megan C. et al., 2019). TEs’ proximity to effector genes interfere with their structure and expression by ORF disruption, promoter insertion, spread of repeat-induced point mutations (RIPs), and epigenetic silencing (Sánchez-Vallet et al., 2018). These processes lead to AVR gene loss of function, structural changes or silencing, which help pathogens escape the recognition of *R* genes. Notably, around 75% of the *Bh* genome comprises TEs and many TE families are actively transcribed (Spanu et al., 2010; Qian et al., 2023). However, information on TE transcription is lacking for individual TE insertion sites. The barley mildew genome, alongside the first chromosome-level *Blumeria* genome from powdery mildew *Blumeria graminis* f.sp. *tritici*, showed that diverse effector clusters were interspersed with massive copies of transposable elements in both species (Frantzeskakis et al., 2018; Müller et al., 2019). Repeat induced point mutation machinery is missing in *Bh* as well as *Bgt* genomes, and this may have caused TE accumulation and genome expansion (Spanu et al., 2010). RNA interference can serve as an alternative mechanism to regulate TE, and long non coding RNAs occuring as TE-antisense transcripts have been identified as potential regulators of TE in *B. hordei* (Qian et al., 2023). However, whether and how TEs also contribute to the innovative potential of *Bh* genomic features was not yet systematically addressed.

In this study, we are interested in the regulatory and structural bases of TEs that lead to the diversity of effector genes and in TEs’ potential role in forming new genes. We applied deep mass spectrometry-based proteomics (MS), long-read and short-read sequencing on both DNA and RNA. Based on these multi-omic resources, we completed new high-quality genome assemblies, gene and TE annotation, and effector prediction to further understand the mechanisms by which TEs impact genes and the genome structure of *B. hordei*. We extended this to identify proteins expressed from supposedly non-coding parts of the genome. Together, this suggests high innovative potential derived from TE in shaping the transcriptional landscape of *B. hordei* and birth of novel protein-encoding genes.

## Results

### Two near telomere-to-telomere genome assemblies of *Blumeria hordei*

To resolve the genome structure of *Bh*, we constructed two high-quality assemblies for the two *Bh* isolates TUM1 and AUS1 using DNA from haploid conidiospores and a combination of PacBio HiFi, Oxford Nanopore, and Hi-C sequencing. The assembly of TUM1, a German *Bh* isolate, has a genome size of 117.2 Mb and comprises 60 scaffolds with a contig N50 value of 10 Mb. Similarly, the assembly of AUS1, a *Bh* isolate collected in Australia, has a genome size of 116.6 Mb and consists of 15 scaffolds with an N50 of 10 Mb (Table 1). BUSCO values for both assemblies are above 98% for core ascomycete genes (OrthoDB v10), indicating that these assemblies are of high quality (Kriventseva et al., 2019). Both assemblies contained 12 scaffolds larger than 1 Mb, as well as a single scaffold of approximately 100 kb that corresponds to the mitochondrial genome. All remaining scaffolds were less than 150 kb. The 12 large scaffolds (>1Mb) were present in a 1-to-1 fashion in both assemblies (Fig. 1, 2). On these 12 scaffolds, we identified 20 and 17 telomeric sequences in the AUS1 and TUM1 assemblies, respectively. In AUS1, 8 out of the 12 large scaffolds carried telomeric sequences at both ends, indicating that they represent complete chromosomes, whereas four scaffolds had telomeric sequences only at one end. In TUM1, 5 of the large scaffolds had telomeres at both ends and 7 scaffolds had telomeric sequences on one end (Fig. 1). In addition, 2 small contigs also contained telomeric sequences in TUM1. In *Bgt*, it was reported that the assembly of chromosome 09 was complicated by the presence of a large tandem array of rDNA (Kunz et al., 2025). Similarly, in both the TUM1 and AUS1 assemblies, we found two large contigs that each carry an rDNA cluster at one end of the scaffold, suggesting that these two scaffolds together may constitute a single chromosome. Together with the Hi-C data, these results indicate that *Bh* has 11 chromosomes. In agreement with this, we identified a single region per chromosome that showed markedly lower GC content, higher TE content and lower or absent gene content (see below), (Fig. 1) that likely represent the centromeres. In summary, we conclude that *Bh* likely has 11 chromosomes.

**Figure 1.**
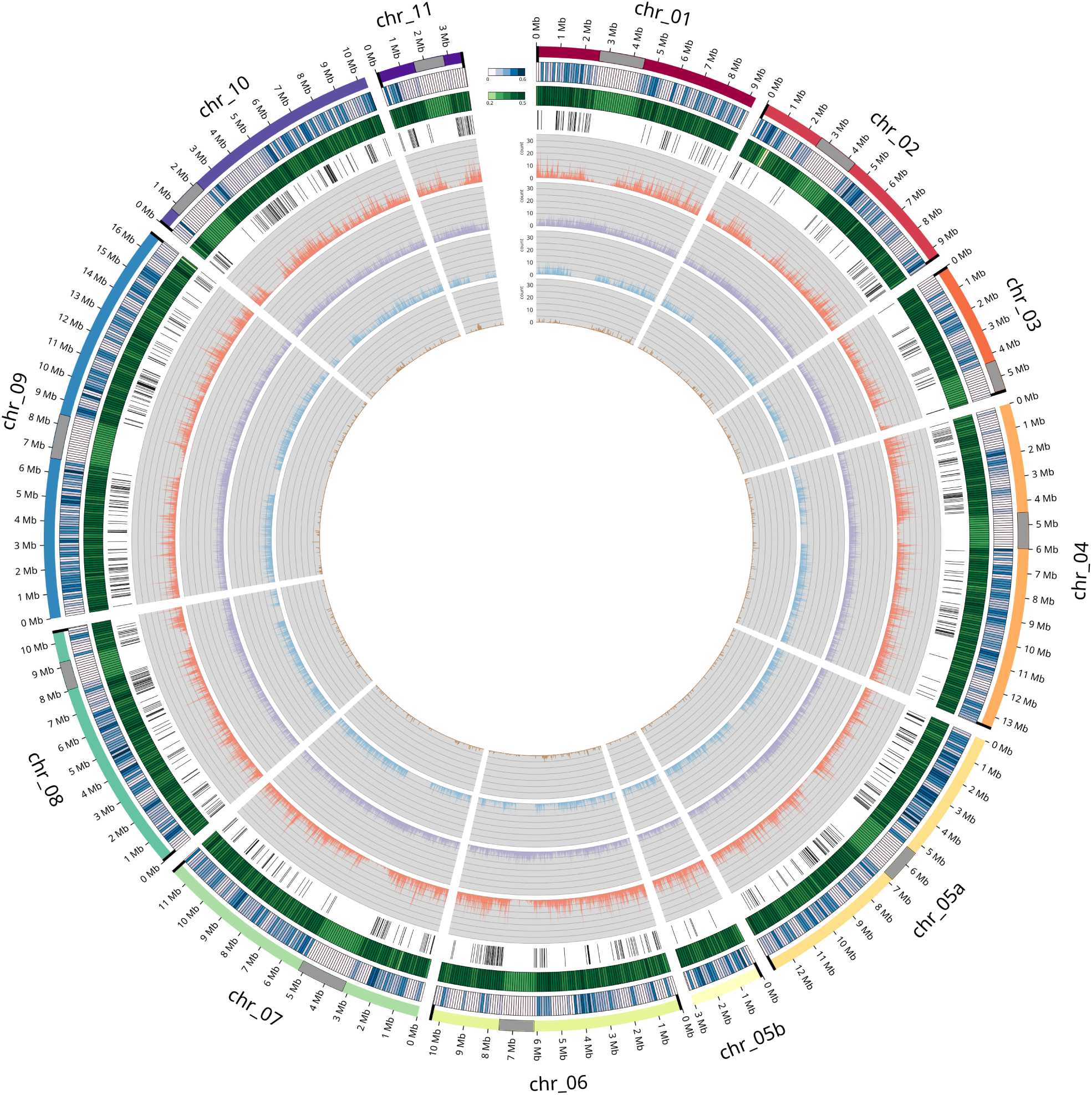
The Circos plot shows the genomic features of the *Blumeria hordei* isolate TUM1. From outer to inner circle: karyotype of 11 chromosomes. Grey regions highlight the centromeric positions of 11 chromosomes. Bold black lines at the ends of chromosomes display assembled telomeres. Gene density is presented in 100 kb windows. GC content in 100 kb windows. Black lines mark the locations of annotated effectors; Line plots show transposable elements (TEs) in 10 kb windows, including long terminal repeats (LTRs), long interspersed nuclear elements (LINEs), short interspersed nuclear elements (SINEs), and terminal inverted repeats (TIRs).

**Figure 2.**
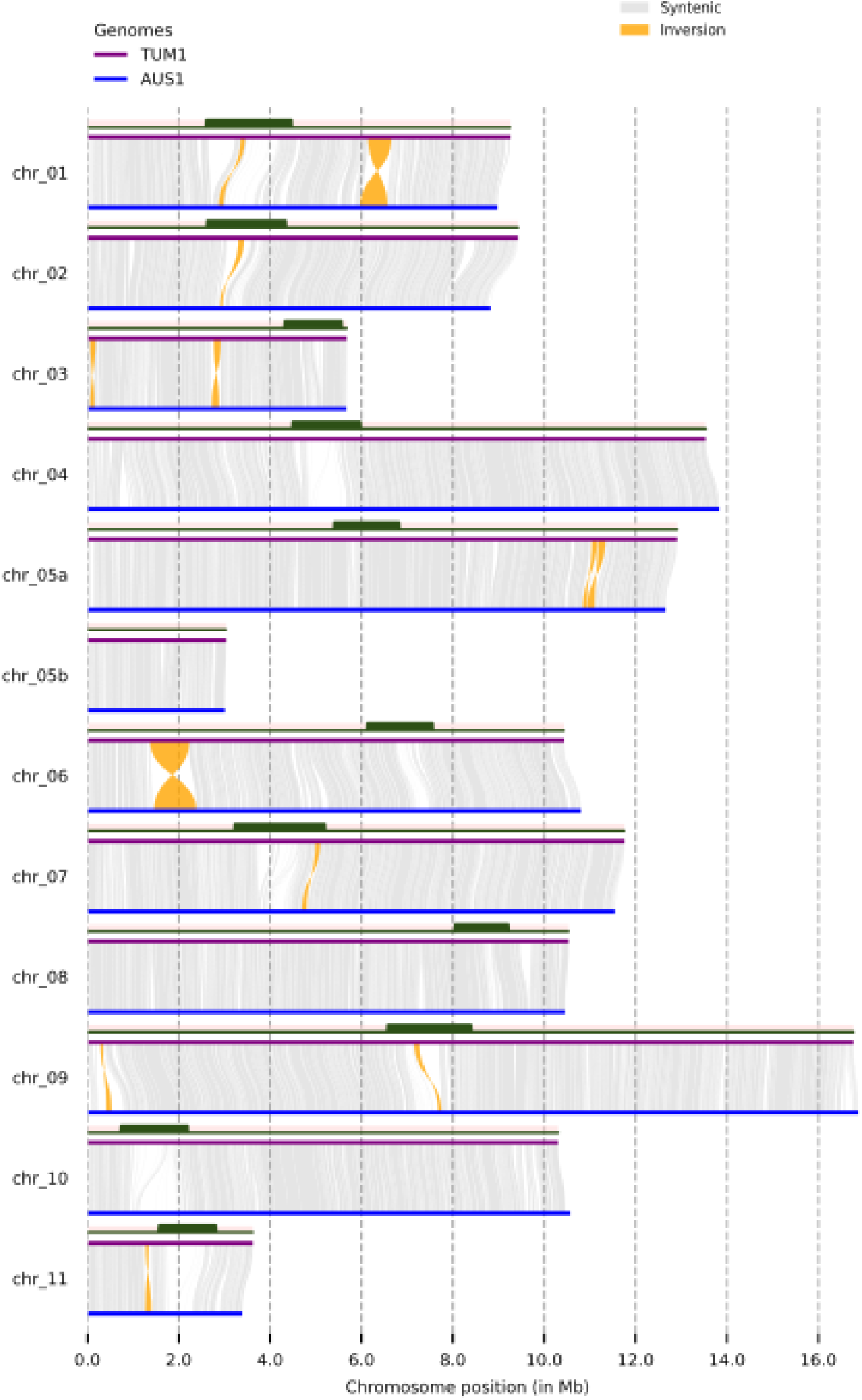
A synteny plot illustrates the structural variation between TUM1 (purple) and AUS1 (blue). Red blocks over red lines represent the positions of centromeres in TUM1. Syntenic regions are gray. Structural variations involved in the analysis include inversions, translocations, and duplications. Structural variations greater than 100 kb were illustrated. Only inversions (yellow) bigger than 100 kb were detected.

**Table 1.** Genome assembly statistics for isolates TUM1 and AUS1.

| **Metric** | **TUM1** | **AUS1** |
| --- | --- | --- |
| Isolate | TUM1 | AUS1 |
| Pseudomolecule number | 12 | 12 |
| Scaffold number | 60 | 15 |
| Telomere number | 19 | 20 |
| Chromosome size (Mbp) | 117.2 | 116.6 |
| Genome size (Mbp) | 118.6 | 116.8 |
| N50 (Mbp) | 10 | 10 |
| BUSCO (ascomycota_odb10) | 98.50% | 98.70% |
| BUSCO (ascomycota_odb12) | 95.30% | 95.40% |

### Genome annotation of *Blumeria hordei*

To annotate *Bh* protein-coding genes in the new genome assembly TUM1, we established a pipeline combining homology-based gene prediction using published *Bh* gene models and *ab initio* predictions supported by transcriptome sequencing and proteomic data. First, we transferred gene models from DH14 (Frantzeskakis et al., 2018) to the new assembly TUM1, comprising 7,077 genes corresponding to DH14 in TUM1. Next, new gene structure predictions were generated with the BRAKER and MAKER pipelines, supported by Illumina RNA-seq dataset of TUM1, and public protein databases from OrthoDB and *Bgt* (Müller et al., 2019), respectively. The transcriptomic evidence was obtained from deep, long- and short-read RNA sequencing. To obtain a time-serial transcriptome capturing genes expressed during different early *Bh* infection stages, we collected TUM1-infected barley epidermal samples at multiple post-inoculation timepoints in triplicate and obtained the corresponding Illumina short-read RNA-seq dataset. The dataset encompassed conidia germination (1 hpi), appressorium development (10 hpi), haustorium formation (20 hpi and 30 hpi), and haustorium maturation (40 hpi and 50 hpi). The rates of aligned short *Bh* reads to TUM1 ranged from 34.07% to 68.80%. Additionally, we used RNA from the 10, 20, 30, and 50 hpi timepoints for full-length cDNA sequencing using PacBio SMRT sequencing technology (PacBio Iso-Seq). We used the standard Iso-Seq workflow to generate full-length, non-concatemeric reads (FLNCs), then identified transcript models from the four Iso-Seq datasets to represent gene isoforms and predicted ORFs. To integrate all homology-based and *ab initio* predictions, we built a pipeline to benchmark different predictions and to select the optimized gene models using the *InGenAnnot* suite, which was originally published for gene annotation of *Zymoseptoria tritici* (Lapalu et al., 2025). Finally, we manually curated gene models that were poorly supported by long- and short-read transcriptome data. In total, our new prediction identified 7806 protein-coding genes. In addition, we used Iso-Seq data from all four time points to define long non-coding RNAs (lncRNAs). We defined lncRNAs as transcripts longer than 200 bp that originated from the anti-sense strand of protein-coding genes, or the anti-sense strand of annotated TEs, or intergenic regions, all lacking SignalP or BLASTp evidence, in accordance with previous studies on lncRNA from *Bh* and *Z. tritici* (Qian et al., 2023; Lapalu et al., 2025). Applying these criteria, we identified 3,270 long non-coding RNA (lncRNA) genes in TUM1. In addition, we used MFannot to identify 28 mitochondrial transfer RNA (tRNA) or ribosomal RNA (rRNA) genes. We compared the TUM1 gene annotation to the DH14 gene annotation. After benchmarking, the TUM1 annotation retained 6,962 gene models from the DH14 gene annotation, along with 844 novel, previously unannotated gene models. To characterize the completeness of our gene annotation, we performed BUSCO analysis on TUM1 protein-coding genes using the latest OrthoDB v12 Ascomycota dataset, and more than 95% were covered (Tegenfeldt et al., 2025). The TUM1 gene models were transferred to the AUS1 assembly, yielding 7,745 corresponding genes corresponding to a BUSCO value of 95.4%.

### Transposable element annotation

To study the TE dynamics in *Bh*, we aimed at obtaining a genome-wide TE annotation with an accurate consensus TE library, particularly for SINEs. SINEs represent an important TE order with potential functional relevance, as previous studies have shown that effector candidate ROPIP1 originates from *Eg_R1*, a SINE family (Nottensteiner et al., 2018). However, SINEs in *Bh* were underrepresented in the latest TE library from Qian et al. compared to those in *Bgt* (Qian et al., 2023; Müller et al., 2019) for which 11 SINE families are described. We therefore manually curated *Bh* SINEs based on the *Bgt* SINE elements in the TREP database and the well-characterized *Eg-R1* and *Egh24* elements. In addition, we manually checked and reclassified the superfamily assignment of TEs in the *Bh*-specific TE library from Qian *et al*., 2023 based on domain identifications and the homology of consensus sequences to TREP and Repbase (Wicker et al., 2002; Jurka et al., 2005; Qian et al., 2023). The new TE library comprises 352 consensus TE sequences, of which we identified 213 LTRs, 99 LINEs, 22 SINEs, 2 TIRs elements, and 16 fragmented retrotransposons. Our comprehensive annotation of TEs using the curated TE library revealed that 73.75% of the TUM1 genome and 73.85% of the AUS1 genome were composed of interspersed repetitive elements, similar to the TE proportion reported for DH14 (Qian et al., 2023). Based on this annotation, class I transposons constituted the majority of the annotated transposable element (TE) length in the *Bh* TUM1 genome, comprising 31.72% LTRs, 29.86% LINEs, and 10.53% SINEs. The only Class II transposon order detected was TIRs, which accounted for 0.41% of the genome. A similar proportion of individual TE orders was observed in the *Bh* AUS1 assembly. Genome-wide TE annotation of TUM1 showed that the indivdiual TEs orders were distributed throughout the genome of TUM1, but centromeric regions were enriched for LINEs and depleted for other TEs (Fig. 1).

### Effector prediction of *Blumeria hordei*

Next, we aimed to identify putative effector genes from the TUM1 gene annotation. To do so, we performed proteome-wide prediction of signalp peptide, transmembrane domains and function prediction for all predicted proteins of TUM1. Subsequently, we performed Markov clustering of the predicted TUM1 proteome with the 531 CSEP from the DH14 reference proteome defined in Sabelleck et al. (Sabelleck et al., 2025) and 844 predicted effectors from *Bgt* (Müller et al., 2019), resulting in the identification of 494 clusters. We define TUM1 proteins as effectors if they meet one of the following critria: (1) Proteins clustered with *Bh* CSEPs (Pedersen et al., 2012; Sabelleck et al., 2025) that were not annotated as conserved functional protein. (2) Proteins clustered with *Bgt* effectors that contain a signal peptide without a transmembrane domain and are not annotated as conserved functional proteins. In this category, if proteins only clustered with *Bgt* but not any *Bh* CSEPs, dit not contain a signal peptide or EffectorP prediction, we manually curated these clusters to make sure that the *Bgt* effectors included in the clusters are true and these TUM1 proteins were not conserved functional proteins. (3) Proteins with a predicted signal peptide, no predicted transmembrane domain and predicted as effectors by EffectorP. Using these criteria, we predicted a total of 919 candidate effectors in TUM1.

Among these 919 candidates effectors, the largest group, namely 408 proteins, consists of predicted effectors present in clusters with both *Bh* CSEPs and *Bgt* effectors (Fig. 7. In addition, we identified 172 candidate effectors that clustered only with *Bh* CSEPs and 162 candidate effectors that clustered with predicted *Bgt* effectors but not *Bh* CSEPs. In our annotation, 580 proteins were predicted to be CSEPs and mapping to 66 CSEP families defined by Pedersen et al. (2012); 438 of these showed high sequence identity to previously reported CSEPs (Pedersen et al., 2012). From the predicted effectorome, 570 genes could be classified into 125 *Bgt* effector families (Müller et al., 2019). Using SignalP, we predicted that 86% (790 of 919) of the predicted effectors encode signal peptides. The prediction of effector candidates without signal peptide is consistent with recent studies from *Bh* and *Bgt*, which identified virulence and avirulence effectors lacking clear signal peptide prediction (Kunz et al., 2026; Weiß et al., 2025). These effectors are part of large, expanded effector families (Kunz et al., 2026; Seong and Krasileva, 2023). We then compared our effector prediction in *Bh* TUM1 with the prediction from *Bh* DH14. We performed BLASTp analyses using DH14 protein sequences as query sequences against the TUM1 effector set. This analysis mapped 817 DH14 genes to 826 TUM1 effectors. Thus, 93 of the 919 identified TUM1 candidate effectors were absent from the DH14 annotation and were identified due to our new annotation strategy.

### Comparative genomics revealed conserved genomic structures between two *Blumeria* species

Previous studies on *B. graminis* have shown that it also possesses 11 chromosomes (Müller et al., 2019). We were therefore interested in how well the chromosomal structures are conserved between *Bh* and *Bgt*. To do so, we compared genome structure based on gene synteny of conserved genes between the two species. Using the OrthoFinder suite, we identified 5309 single-copy orthologous genes shared between *Bh* and *Bgt*, corresponding to 68% and 68.5% of the predicted gene complements of TUM1 and AUS1, respectively. This analysis revealed overall high conservation of gene synteny between *Bgt* and *Bh* (Fig. S1, Fig. S2). Notably, for seven of the eleven *Bh* chromosomes (Chr01, Chr03, Chr06, Chr07, Chr08, Chr10, Chr11), more than 95% of the conserved genes per chromosome were located on a single corresponding chromosome in *Bgt*, indicating that these seven chromosomes are present in a 1-to-1 fashion in both species. In contrast, we detected two large-scale rearrangements between Chr02 and Chr04, occurring on the chromosome arms, as well as a centromeric rearrangement between Chr05 and Chr09. In addition to these inter-chromosomal rearrangements, we also identified a large inversion in the central region of Chr06, affecting approximately 4.8 Mb compared to the corresponding Chr06 of *Bgt* (Fig. S3, S4). To facilitate comparative studies between *Bgt* and *Bh*, we named the *Bh* chromosomes according to their corresponding *Bgt* chromosome numbers whenever possible. The unresolved *Bh* chromosome 05 was remained subdivided into Chr05a and Chr05b. In summary, this analysis suggests highly conserved gene synteny despite the sequence divergence in the intergenic space between the two species.

### Large-scale rearrangements between geographically distant *Bh* isolates

To better understand the sequence divergence between *Bh* strains, we compared the genome sequences of the TUM1 and AUS1 isolates collected from geographically distant regions. We performed a genome alignment using the 12 assembled chromosome-level scaffolds of TUM1 and AUS1. We observed that 98.4%-99.4% of the genomes were aligned. From this alignment, we identified 4,155 syntenic regions between the two assemblies, accounting for 81.97% and 83.01% of the TUM1 and AUS1 assemblies, respectively. The non-syntenic regions were enriched in centromeres (Fig. 2). Structural variations (SVs) accounted for 20.08-21.24 Mb in two assemblies, including 4.89-4.93 Mb inversions, 6.01-6.10 Mb translocations, and 9.19-10.22 Mb duplications (Table. S2). Among structural variants, inversions were the most abundant and were primarily observed between homologous chromosomes. These included a 0.85 Mb inversion located on chromosome 6 and another 0.51 Mb inversion on chromosome 1. For both inversions, the break points at TUM1 were inspected and confirmed in our long read data and covered or surrounded by retrotransposons. To determine whether the inversion originated in AUS1 or TUM1, we examined the synteny of the genes located in the inverted region relative to the gene order in *Bgt* (Fig. S3, S4, S5, S6). For both inversions, the gene order within the inverted segment of TUM1 matched that observed in *Bgt*, whereas the gene order in AUS1 was inverted relative to *Bgt*. This indicates that the TUM1 configuration represents the ancestral gene orientation and suggests that the inversions has occurred in the AUS1 isolate after separation of the two species.

### Iso-Seq identified transcripts from TE and genes

Next, we aimed at understanding the diversity of the transcriptional landscape of the *Bh* genome, including the expression of genes and individual TEs. Iso-Seq data is highly sensitive for detecting transcriptional activity, whereas short-read sequencing is more suitable for accurate transcript quantification. We therefore used long-read RNA-seq data to identify full-length transcripts and short-read RNA-seq data of corresponding time points to quantify their expression levels. We filtered representative transcript models to retain isoforms lacking evidence of intra-priming and supported by at least 10% of all reads assigned to the corresponding gene locus. Subsequently we assessed the quality of these transcript models using SQANTI3 to identify both conserved and novel transcript isoforms (Pardo-Palacios et al., 2024). After filtering, we detected 27,782-34,490 transcript models across the four time point-specific Iso-Seq datasets. We classified isoforms according to their level of agreement with the TUM1 reference gene models. Based on the SQANTI3 classification scheme, we assigned isoforms to the following categories: isoforms that aligned to reference genes with full splice match (FSM), isoforms that have the same splice junctions as the reference genes but incomplete [incomplete splice match (ISM)], isoforms that share splice junctions but contain novel combinations [novel in catalog (NIC)], and isoforms contain novel splice junctions [novel not in catalog (NNC)], isoforms originating from the antisense strand of reference genes, and intergenic transcripts. A total of 14,907–16,973 (49.21%-53.66%) isoforms were associated with genes and 10,594-14,659 (38.13%-42.50%) of isoforms originated from the intergenic space (Fig. 4). These isoforms in genic regions corresponded to 6,356–7,057 genes. A total of 6,153 (78.82% of all genes) genes were present in all four Iso-Seq datasets, whereas an additional 1,193 genes showed time-point specific expression. A total of 809 effectors were identified as transcribed in Iso-Seq datasets, increasing over time from 511 at 10 hpi to 754 at 50 hpi. A total of 479 effector genes were detected in all four Iso-Seq datasets (Fig. 5). Despite the overall increase, some effectors were detected exclusively at early or late time points. On average, we detected 2.59 isoforms per gene locus across all time points, whereas effector genes have an average of 2.02 isoforms. Among the genic isoforms detected, the largest proportion, namely 31.21%-34.65% of total transcripts, belong to novel splice junctions of annotated genes (NNC) while fully matched isoforms corresponding to genes (FSM) only accounted for 13.3%-13.8% of total isoforms in datasets. Notably, more than 85% of isoforms containing canonical splice junctions (FSM, ISM, and NIC) were supported by short-read sequencing data. In comparison, fewer NNC isoforms were evidenced by short-read RNA-seq suggesting that some of them are rare transcript isoforms. Yet, we detected short-read RNA-seq support for 55.55% of these NNC isoforms, supporting the presence of novel alternative transcript isoform from genes.

**Figure 3.**
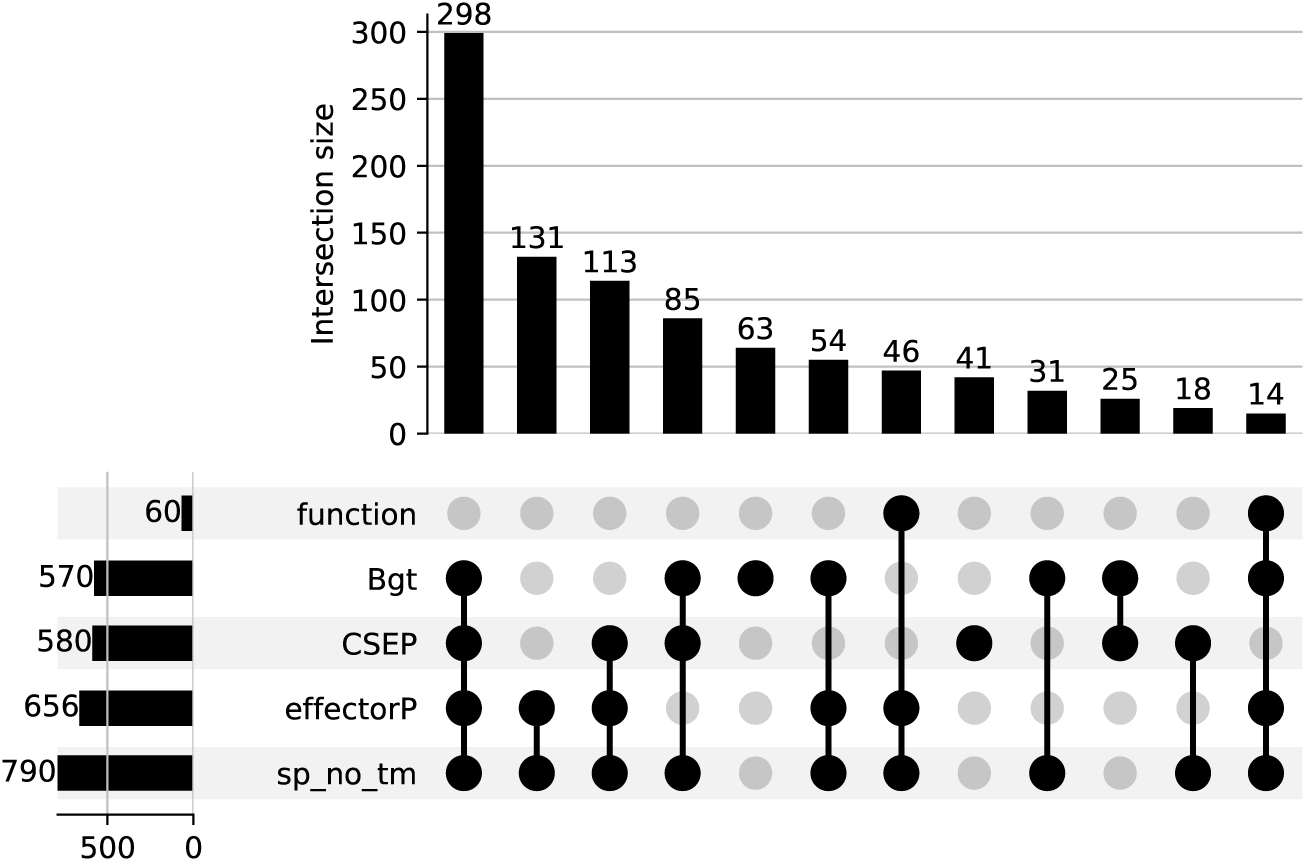
The UpSet plot shows the number of predicted effector supported by different combinations of evidence. Function: Effector is predicted to be a functional protein by PANTHER (Thomas et al., 2022). Bgt: Effector is clustered with predicted *Bgt* effectors (Müller et al., 2019). CSEP: Effector is clustered with Candidate Secreted Effector Proteins (CSEP) (Pedersen et al., 2012). EffectorP: Predicted effector from EffectorP 3.0 (Sperschneider and Dodds, 2022). sp_no_tm: Effector has signal peptide without a transmembrane domain (Teufel et al., 2022; Hallgren et al., 2022).

**Figure 4.**
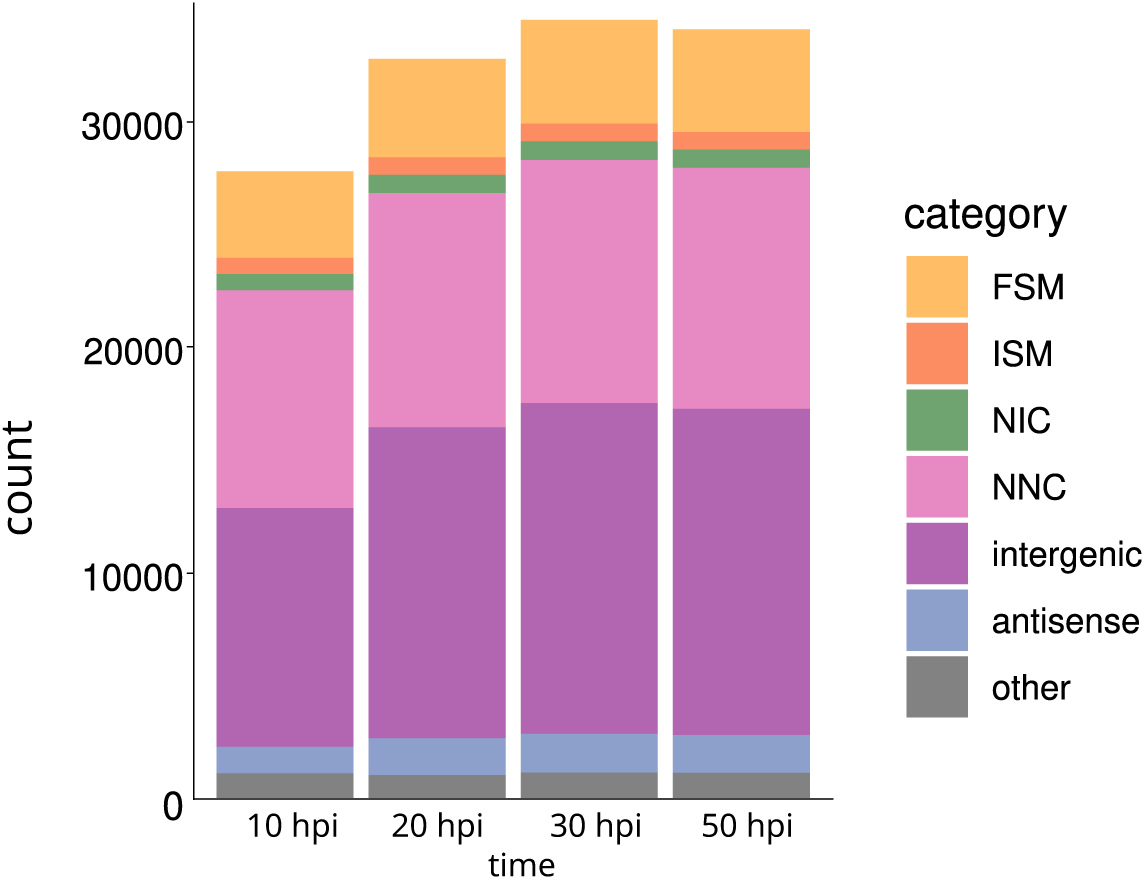
The overview of identified isoforms from four Iso-Seq libraries by the output of SQANTI3. FSM (Full Splice Match): isoforms have the identical number of exons and splice junctions as annotated genes. ISM (Incomplete Splice Match): isoforms have fewer 5’ exons than the annotated genes, but aligned splice junctions. NIC (Novel In Catalog): isoforms only use a combination of known splice sites of annotated genes. NNC (Novel Not in Catalog): isoforms have at least one novel splice site. Intergenic: isoforms transcribed from intergenic regions. Antisense: Isoforms transcribed from genic regions but represent the antisense strand. Other: transcripts at genic introns or transcripts run through introns and exons, which can be artifacts.

**Figure 5.**
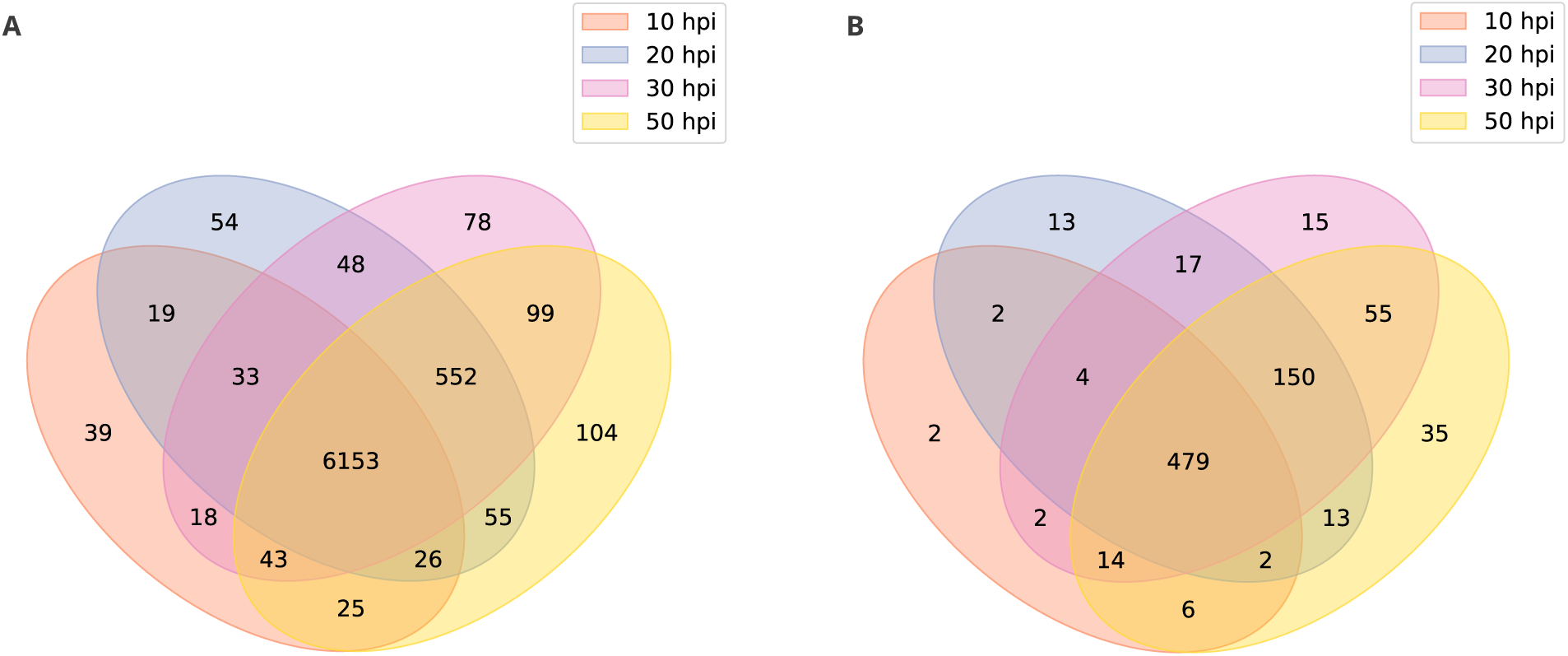
A. The Venn diagram shows the number of all annotated genes identified in the Iso-Seq datasets and classified by SQANTI3. B. The Venn diagram shows the number of effector genes identified in the Iso-Seq datasets and classified by SQANTI3.

Next, we aimed at characterizing general TE transcription. Our long read DNA and RNA-seq data together allowed for insertion site-specific analysis of transcripts from individual TEs. Based on Iso-Seq transcript data, we hence assessed the percentage of TE loci in the genome that produce transcripts. We found that 31.71% of TE loci are transcribed. 36.16%-47.79% of the 29,964 annotated genomic SINE accessions were transcribed. Thus SINEs were the most transcriptionally active TE order despite lacking TE protein-encoding capactiy. TIR elements account for less than 0.5% of the genomic TE annotation, but also exhibit active transcription, which increased from 29.91% at 10 hpi to 38.89% at 50 hpi. LINEs and LTRs exhibited a stable proportion of transcriptionally active TEs over time, ranging from 27.18% to 32.75% (Fig. 6). We then assessed the contribution of these transcribed TEs to the *Bh* transcriptome using two complementary measures: their contribution to the cumulative length of unique transcripts and their abundance in the short read RNA-seq transcriptome. Among the unique isoforms identified across the four datasets, LTRs and LINEs together accounted for 28.64%–30.75% of the total isoform length at the four time points, whereas SINEs contributed 7.98%–9.98% in each dataset (Fig. 6). TIRs showed an extremely low contribution to total isoform length (<0.5%), reflecting their lower frequency in the *Bh* genome. We then used the proportion of high-quality short RNA reads from Illumina sequencing that mapped to genomic TEs to estimate the transcript abundance of TEs. At different time points of the four isoform datasets, retrotransposons (LINEs, LTRs, and SINEs) cumulatively accounted for 5.19%-5.74% of total reads mapped to the genome (Table S1). TIRs took only a low proportion of the transcriptome (0.01%-0.02%). Among the retrotoransposon, 2.08%-2.29% TE orginated RNA reads originated from LTR elements. Despite their short sequence lengths, SINEs exhibit an average of 1.83% mapped reads across four time points, which is comparable to LINEs (1.40%). Together with their high proportion in unique isoforms, the read quantification suggested TEs only took minimal abundance of *Bh* transcriptome, but are important sources of transcript diversity.

**Figure 6.**
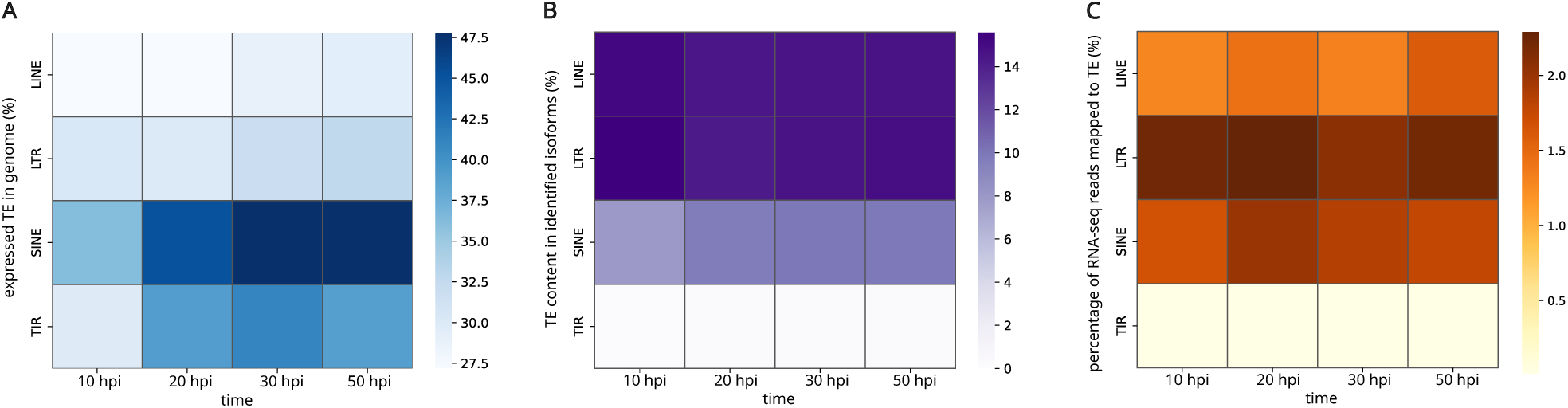
A. The percentages of expressed genomic TEs within each order were calculated by dividing the number of TEs expressed in each order in the Iso-Seq dataset by the total number of annotated TEs belonging to that order. B. The percentages of TE content in different Iso-Seq datasets were calculated as the cumulative TE-masked length relative to the total transcriptome length. C. The percentages of RNA-reads mapped to TEs in different Iso-Seq datasets. The percentages of RNA-seq reads mapped to TE was quantified as the fraction of reads mapped to TEs relative to the total number of reads aligned to the genome.

To understand the origin of actively transcribed TEs, we analysed the genomic TE distribution. We quantified the distance from the border of each TE to its nearest annotated gene and identified TEs located in gene-proximal regions (Fig. 7). On average, SINEs had the shortest distances to genes, whereas LTRs had the longest. In terms of TE–gene distances, significant differences were detected between SINEs and the other three TE orders (*p <* 0.01). Similar patterns were also observed between TIRs and the other orders (*p <* 0.01), whereas no significant difference was detected between LTRs and LINEs. In the flanking regions of genes, SINEs, LTRs, and LINEs were enriched within 8 kb up- and downstream of genes, but SINEs were particularly accumulated within the 2-kb flanking regions (Fig. 8). We detected 4,008 annotated TEs that overlapped genic regions by more than 50 bp across 1,893 genes, including 439 candidate effector genes. In total, these TEs overlapping genic regions accounted for up to 4.06% of total TE accessions across the genome. In genic regions, we observed 3,755 TEs localized in untranslated regions (UTRs). Notably, TE insertions were more frequent in 3*^′^* UTRs (2,588) than in 5*^′^* UTRs (1,167), whereas insertions within ORFs were comparatively rare (253). SINEs again displayed distinctive patterns compared to other retrotransposons: SINEs accounted for 39.01% of TEs in UTRs, whereas LINEs together with LTRs accounted for 58.26%. We defined TEs reaching at least 90% of the length of the consensus TE sequence as intact TEs. Using this criterion, we detected 378 intact TEs within UTRs, of which 334 (88.36%) belonged to SINEs, suggesting that these insertions may be evolutionarily younger or maintained.

**Figure 7.**
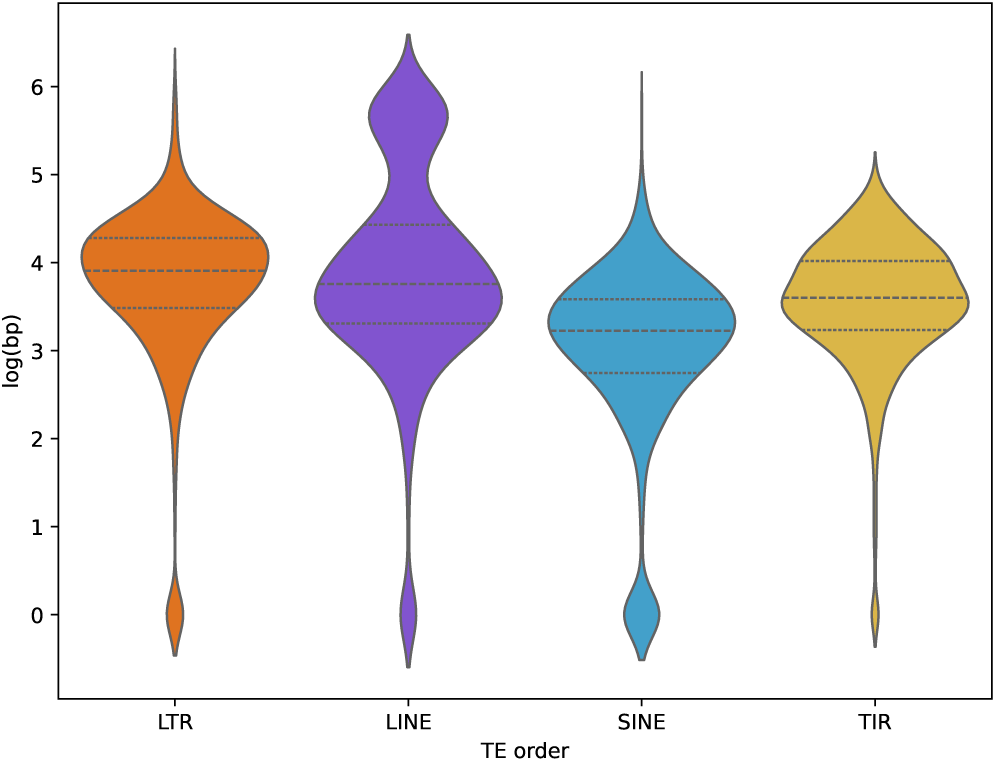
The violin plot depicts distributions of genomic TEs’ distances (bp, log10-transformed) to their closest genes. The dashed lines marked quartiles of TE orders.

**Figure 8.**
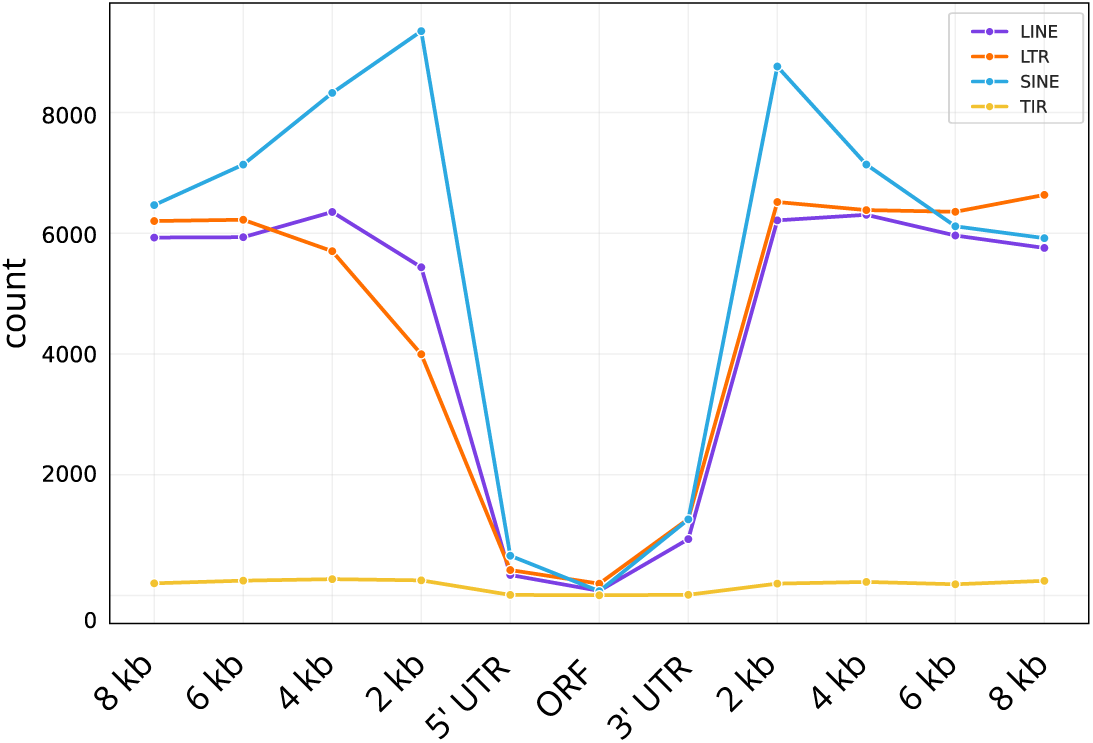
The line plot indicated genomic TE counts inserting into genes and their flanking regions, which included ORFs, 5’ and 3’ untranslated regions (3’ UTR and 5’ UTR), and 0 - 8 kb up- and downstream of the genes.

Transcripts containing TE segments, either intact or fragmented, exhibited two distinct transcriptional modes: autonomous TE transcription and co-transcription with host genes (Fig. 9). For example, we identified 1239–1426 gene–TE chimeric transcripts involving genes and neighboring SINEs, representing the most abundant class of gene-TE chimeric transcripts. In addition, SINEs were frequently transcribed independently, accounting for 416 transcripts at 10 hpi and increasing to 972–1.019 transcripts at later time points. We next compared the expression levels of these transcript classes, quantified as transcripts per million (TPM) based on short-read RNA-seq data. Significant differences in expression were observed between gene transcripts and TE transcripts, as well as between gene transcripts and gene–TE chimeric transcripts (Fig. 10). Across all four Iso-Seq time points, gene transcripts exhibited the highest expression levels, followed by gene–TE chimeric transcripts, whereas TE-only transcripts consistently showed the lowest expression (Fig. 10). Data show a high frequency of transcriptional activity from individual TE loci. TE often insert close to or even into the body of *Bh* genes, the latter mostly into UTRs. Most TE or TE-gene chimera show lower transcript levels when compared to other *Bh* genes. However, individual TE and chimeric TE-gene transcripts reach high transcript levels.

**Figure 9.**
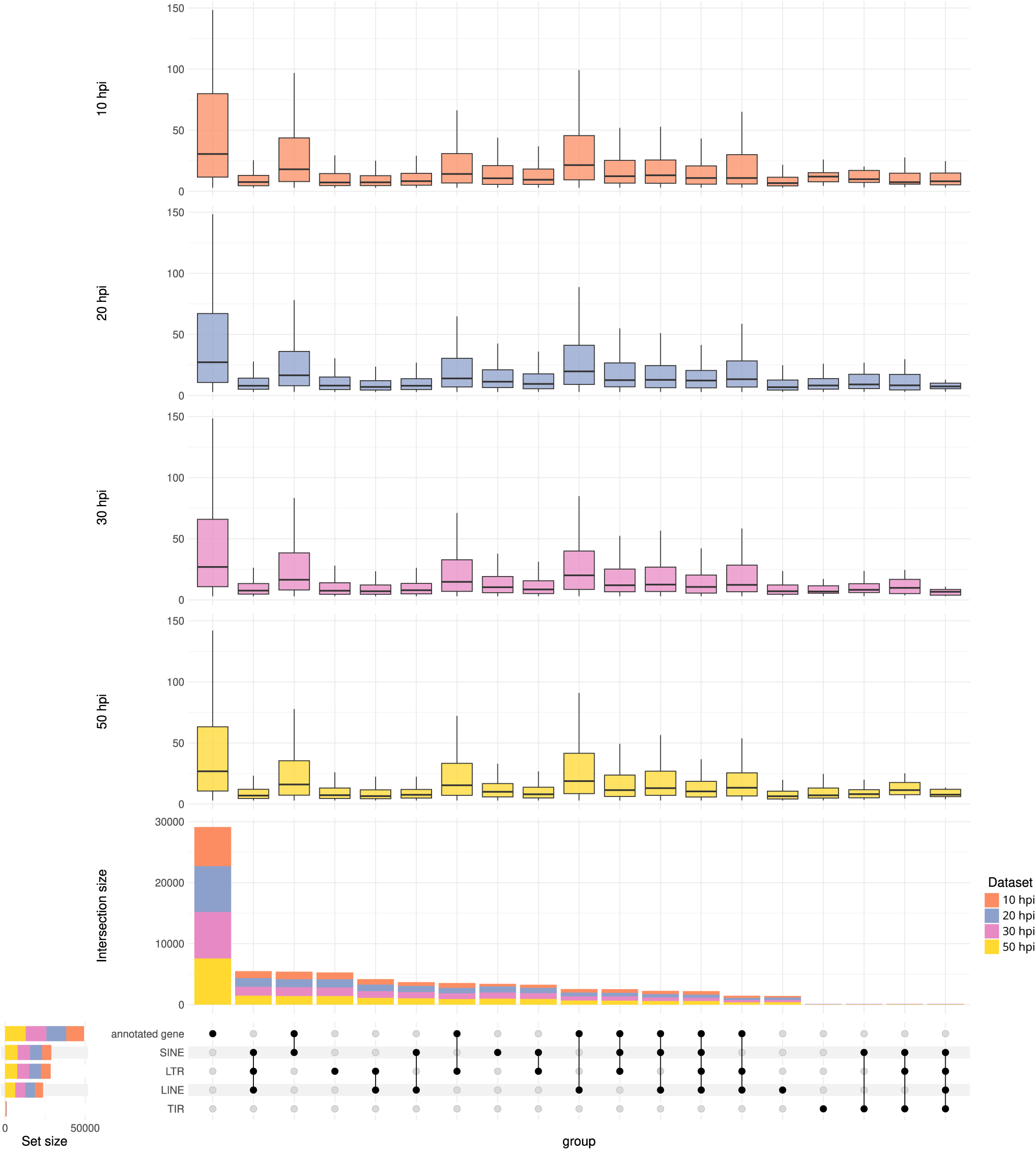
The UpSet plot shows the associated genes and TEs detected in the Iso-Seq datasets. The vertical bar plot illustrated the number of unique transcripts that consist of genes and TEs at different time points. The horizontal bar plot showed the occurrence of TEs/genes at different time points. The box plots show the expression level of transcripts, quantified with Illumina RNA-seq reads and normalized by transcripts per million (TPM). hpi: hour post inoculation.

**Figure 10.**
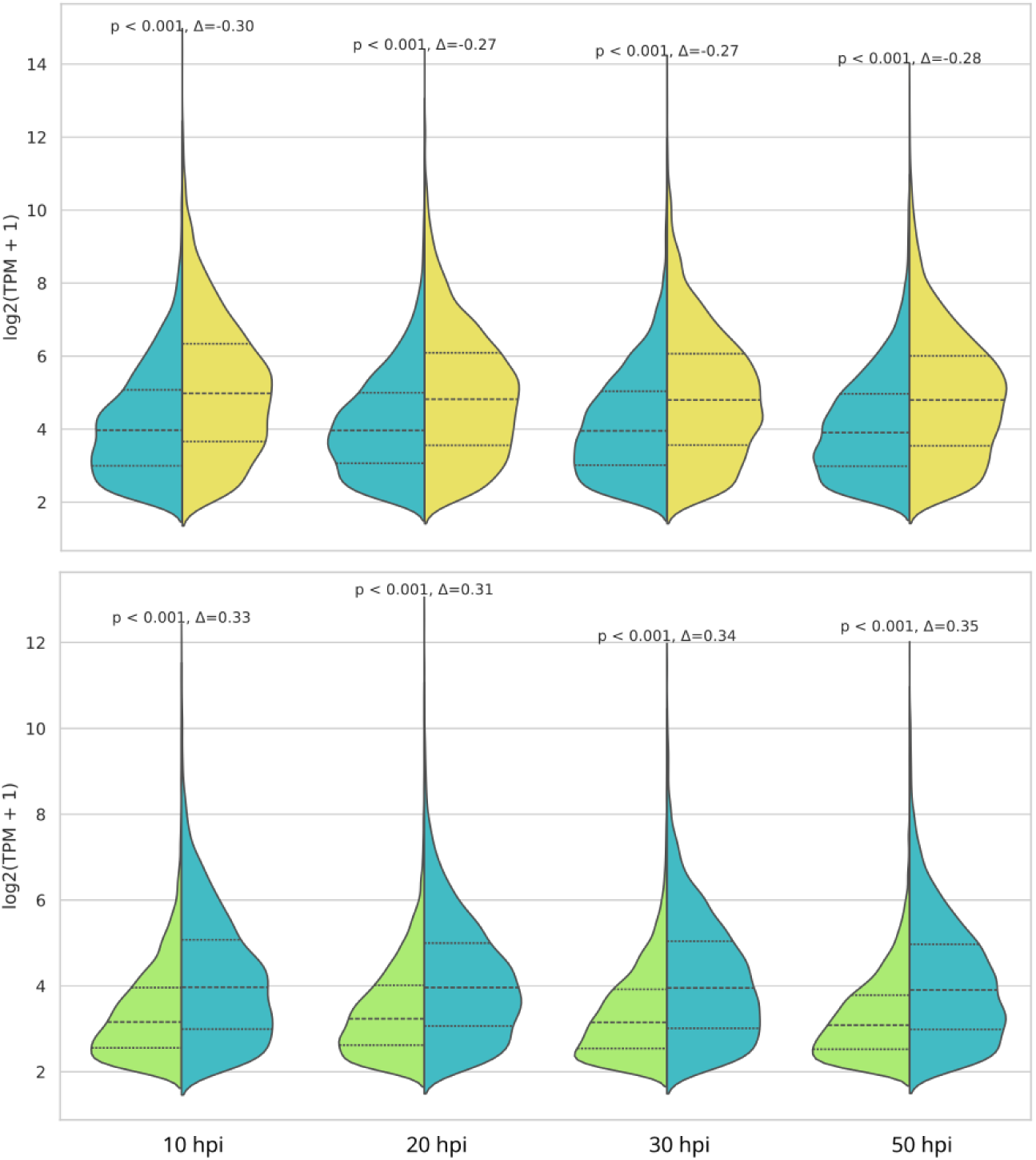
The violin plots compare transcript expression levels within each Iso-Seq library. Lake blue depicts transcripts that consist of both genes and TEs. Lemon indicates transcripts transcribed solely from genes, while green represents transcripts derived exclusively from TEs. Expression levels were quantified as average log2-transformed transcripts per million (TPM). Short RNA reads from the same infection time point were mapped to the reference genome, and TPM values were estimated using StringTie2 with transcript models from Iso-Seq as reference. The median TPM across replicates was then calculated for each transcript. P-values were computed with the Mann-Whitney U test, and Δ indicated Cliff’s Delta, showing the magnitude of differences.

### Applying proteogenomic approach to identify novel proteins

Next, we aimed to describe the protein diversity and TE-derived proteins in the proteome of *Bh*. We hypothesized that diversity arises not only from different gene loci but also from structural differences arising from alternative splicing (AS) and protein complexity. We established a sample-specific proteogenomic pipeline to (1) identify coding sequences based on Iso-Seq and (2) characterize the *Bh* proteome with a hybrid reference database of canonical genes and Iso-Seq-derived coding sequences to provide evidence for gene annotation and to identify potential novel proteins. To do so, we established a sample-specific proteogenomic pipeline that bridges PacBio Iso-Seq transcripts with a bottom-up proteomic database. We used deep liquid chromatography–tandem mass spectrometry (LC–MS/MS) to quantify peptides from extracted proteins of barley epidermal samples infected with TUM1 at 50 hpi. Importantly, the LC–MS/MS samples were derived from the same biological material used for Iso-Seq library construction, and proteins from three replicates were pooled after extraction. We measured 32 pools from 96 HPLC-prefractioned peptide samples. We prepared a hybrid database containing coding sequences (CDS) of annotated genes from both *Bh* TUM1 and barley proteins from the cultivar ‘Golden Promise’ (GP), as well as ORFs generated by three-frame translation of the long read RNA Iso-Seq dataset at 50 hpi. We filtered the predicted ORFs to retain sequences at least 25 amino acids in length. When one ORF was fully contained within another, we retained only the longest ORF. Identical ORFs derived from the same gene locus were represented by a single sequence. We then performed peptide-spectrum match (PSM) in MaxQuant (Tyanova et al., 2016) with a hybrid database serving as a reference database. Mass spectrometry data were processed using a deep-learning–enhanced workflow implemented in the software Prosit, with peptide- and protein-level false discovery rates (FDR) controlled at 1%, enabling high-confidence protein identification and quantification (Gessulat et al., 2019; Picciani et al., 2024).

Mass spectrometry (MS) identifies peptides rather than proteins, and individual peptides can map to multiple protein sequences. Consequently, when all peptides detected for one protein also match another protein, proteins lacking unique peptides cannot be distinguished by proteomic analysis. Therefore, MS-based proteomic data are represented as protein groups (PGs), which summarize the unique peptides assigned to each PG. We defined a protein as novel if it met all of the following criteria: (1) it is not predicted in the current TUM1 gene annotation; (2) it is the leading protein within a PG, defined as having the highest number of matching peptides in that PG; and (3) at least one unique peptide from this PG exclusively maps to this protein. To identify proteins derived from *Bh* and to discover novel proteins based on Iso-Seq data, we classified protein groups (PGs) into three categories according to the source of the leading protein, defined as the protein with the highest number of mapped peptides within each PG. Based on this criterion, PGs were classified as TUM1-annotation-dominated PGs, Iso-Seq-dominated PGs from *Bh*, or barley-dominated PGs. TUM1-annotation-dominated PGs and barley-dominated PGs represent protein groups with leading proteins derived from the TUM1 and barley annotations, respectively. Iso-Seq-dominated PGs contain coding sequences predicted by three-frame translation of Iso-Seq transcripts as their leading proteins. This category includes novel proteins derived from alternative isoforms as well as proteins originating from intergenic regions. We filtered the PGs and peptides inferred by Prosit, retaining those supported by at least 2 peptides or by a single peptide that covered at least 20% of the protein sequence length. We ranked the log10-transformed sum intensities of peptides for each PG category, and their distributions were similar and approximately normal (Fig. 11). This indicates a comparable quality of the prediction across the three PG categories. The density distributions of mapped peptide counts and peptide lengths showed consistency for each PG category, suggesting an average mapped peptide count of 3 and an average peptide length of 10 amino acids (Fig. 12). After quality control of the protein dataset, 4,656 proteins from the TUM1 annotation, 10,422 Iso-Seq-derived pathogen proteins, and 10,651 proteins from the Golden Promise gene annotation were represented across the 13,347 protein groups (Table 2).

**Figure 11.**
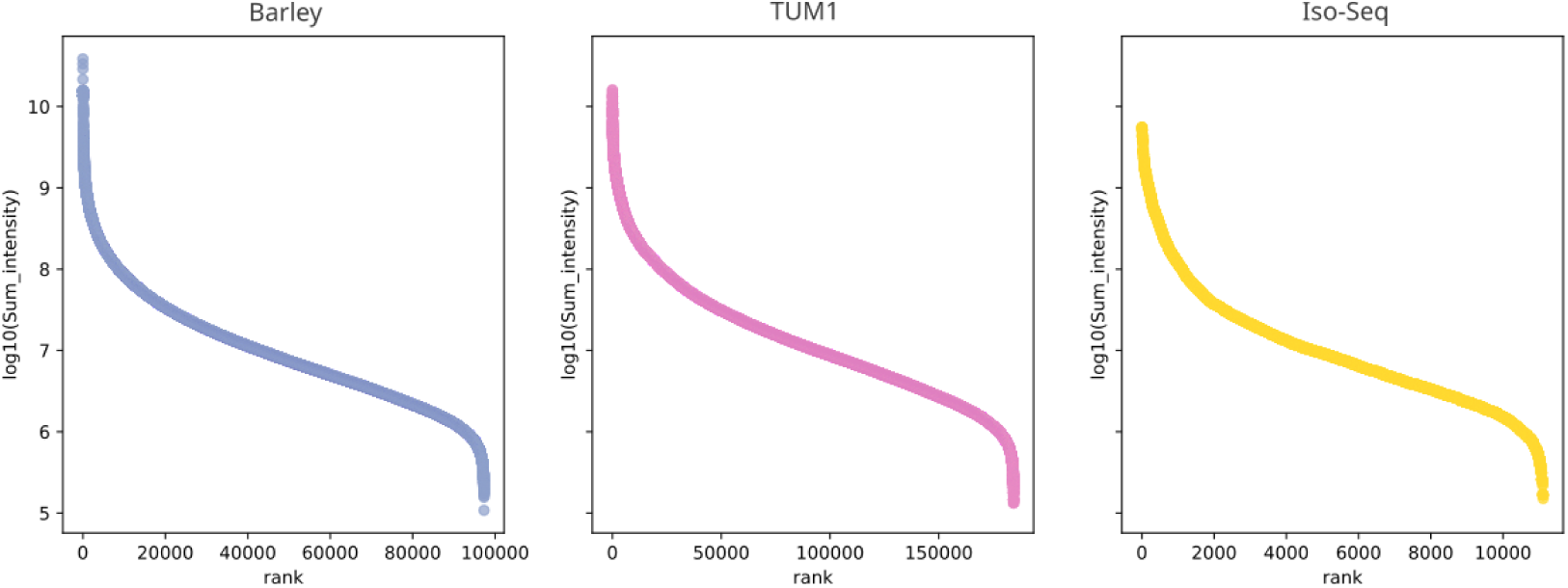
The ranked scatter plot showed the distribution of protein group intensities for sequences detected in each category. Barley: The barley-dominated protein groups. TUM1: The TUM1-annotation-dominated protein groups. Iso-Seq: The Iso-Seq-dominated protein groups.

**Figure 12.**
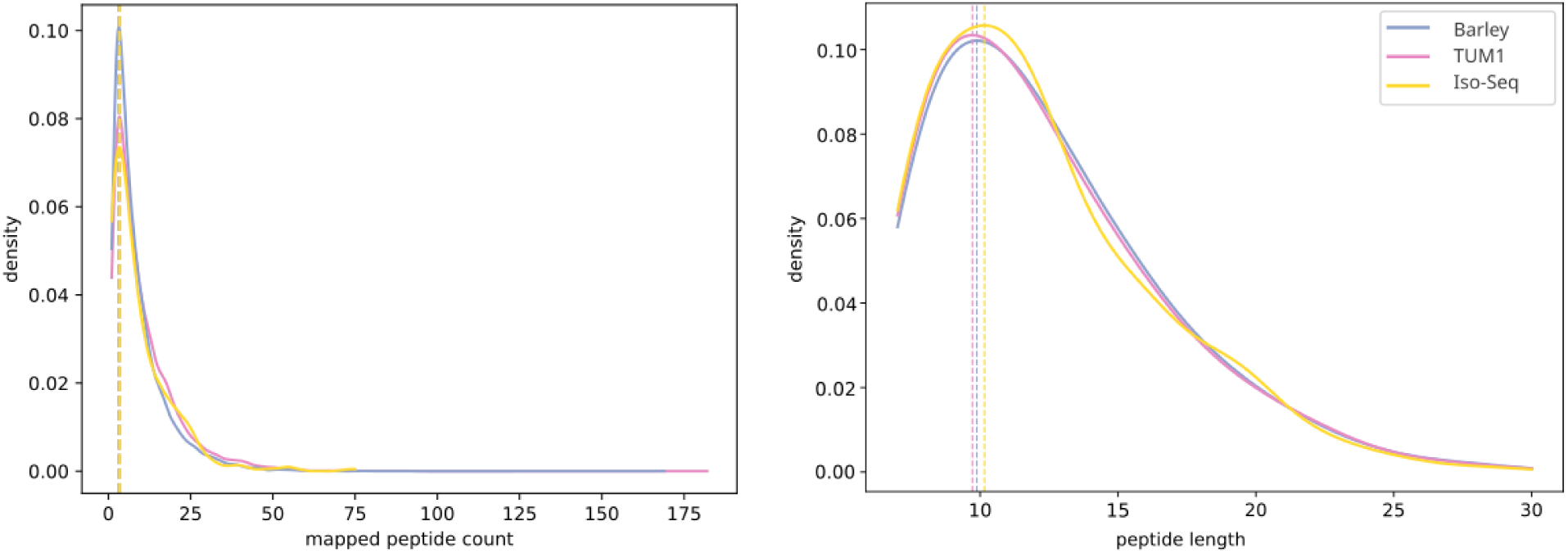
The histograms of peptide counts and peptide lengths for predicted proteins in each category. Barley: The barley-dominated protein groups. TUM1: The TUM1-annotation-dominated protein groups. Iso-Seq: The Iso-Seq-dominated protein groups.

**Table 2.**
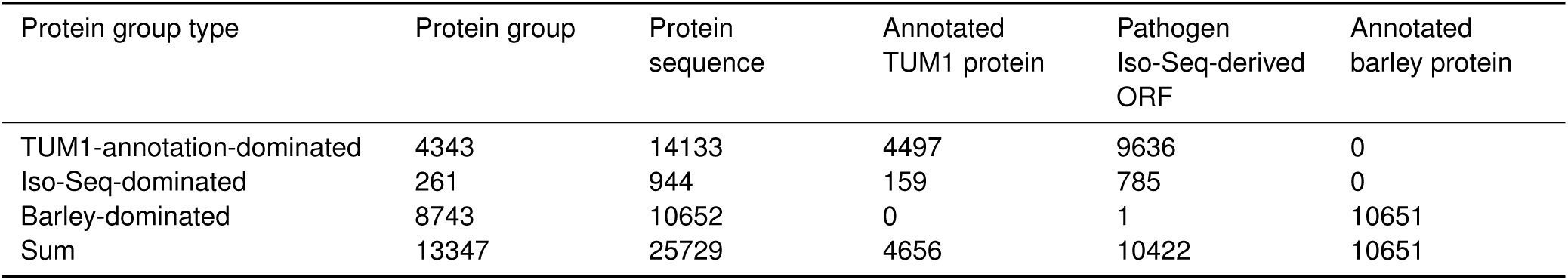
Overview of protein groups from mass spectrometry.

| Protein group type | Protein group | Protein sequence | Annotated TUM1 protein | Pathogen Iso-Seq-derived ORF | Annotated barley protein |
| --- | --- | --- | --- | --- | --- |
| TUM1-annotation-dominated | 4343 | 14133 | 4497 | 9636 | 0 |
| Iso-Seq-dominated | 261 | 944 | 159 | 785 | 0 |
| Barley-dominated | 8743 | 10652 | 0 | 1 | 10651 |
| Sum | 13347 | 25729 | 4656 | 10422 | 10651 |

Using TUM1-dominated protein groups (PGs), we validated the annotation of TUM1 genes: at 50 hpi, short-read RNA-seq data detected expression of 7,420 TUM1 genes, of which translation of 4,643 genes was additionally supported by proteomic evidence (Fig. 13). Among the 849 candidate effectors expressed at 50 hpi according to RNA-seq, 241 were also supported at the protein level. Finally, we assessed the relationship between protein abundance (iBAQ) and transcript levels measured by short-read RNA-seq (TPM) using Spearman’s rank correlation, revealing a moderate positive correlation (0.47, *p <* 0.001) (Fig. 14). We focused on genes that were highly expressed in both datasets, defined as those with values above the median for both iBAQ and TPM. We performed Gene Ontology (GO)–based functional enrichment analysis on the 1,528 genes included within this quadrant. We observed an enrichment of active transcription and translation of housekeeping genes involved in post-transcriptional processes (Fig. 15). Additionally, a total of 63 effectors were identified in this quadrant, 42 of which belong to previsouly described *Bh* and Bgt CSEP families, such as E014/CSEP0064 and E003 (Pedersen et al., 2012; Müller et al., 2019). We found that 11 effectors belong to significant groups in the GO enrichment analysis and 4 effectors, BHTUM1_CHR08056610, BHTUM1_CHR05029880, BHTUM1_CHR04017260 and BHTUM1_CHR09057650, were annotated as ribonuclease-like superfamilies in Pfam and were predicted with the molecular function of RNA binding, indicating that they likely belong to the RNase-like effectors. These four effectors were annotated as CSEPs and conserved across *Bh* and *Bgt*.

**Figure 13.**
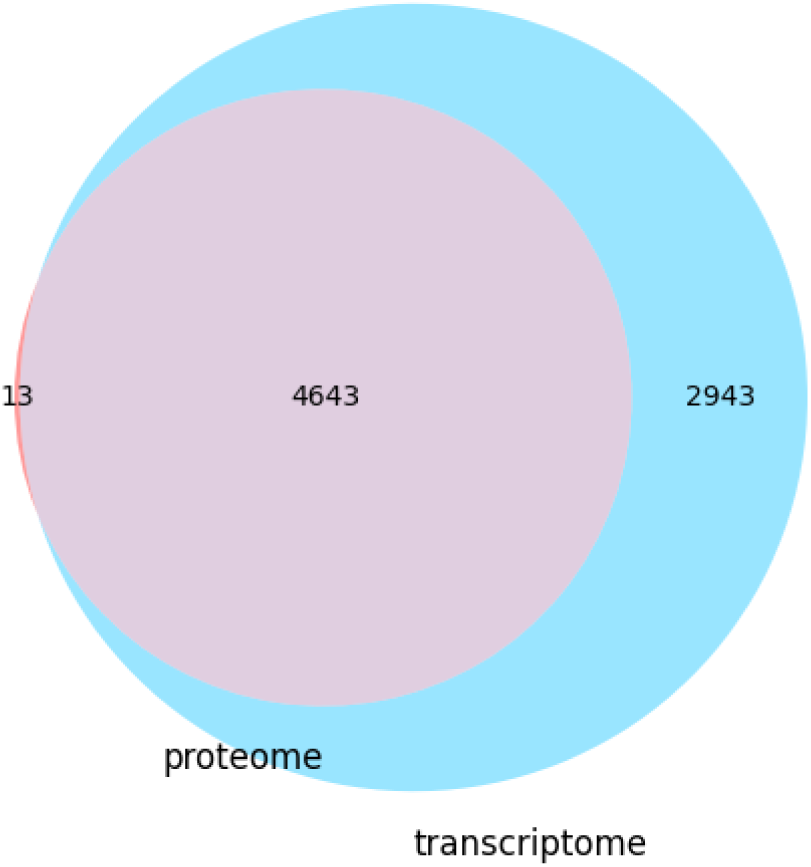
The Venn graph showed annotated *Bh* TUM1 genes present in the proteome and transcriptome. The proteome was represented by Mass Spectrometry (MS) at 50 hours post inculation (hpi). The transcriptome was presented by RNA-seq at 50 hpi.

**Figure 14.**
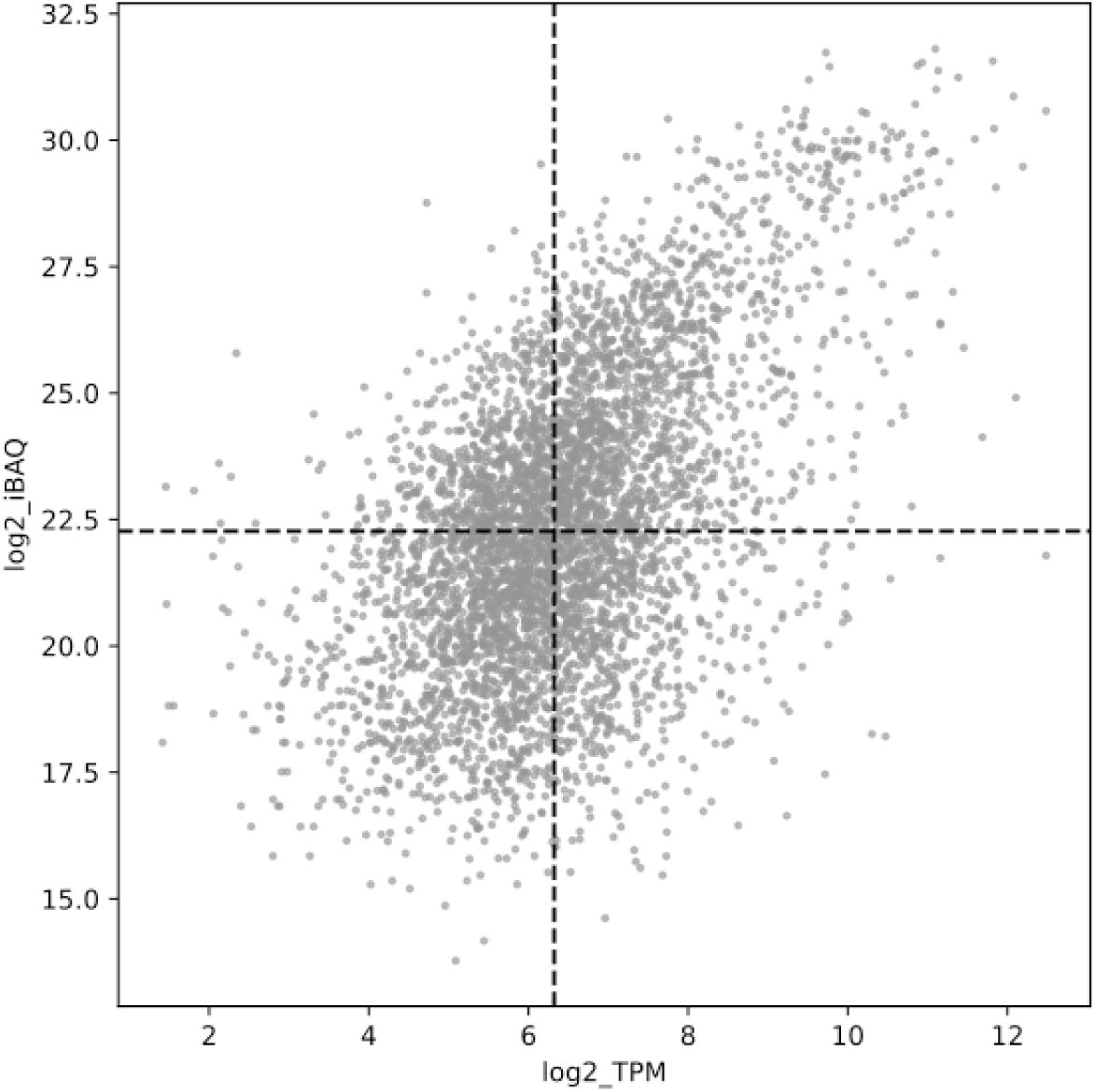
The scatter plot showed the correlation of expression levels of genes supported by Illumina RNA-seq and MS at 50 hpi. log 2_TPM: log 2-transformed Transcripts Per Million (TPM). log 2_iBAQ: log 2-transformed Intensity-Based Absolute Quantification (iBAQ).

**Figure 15.**
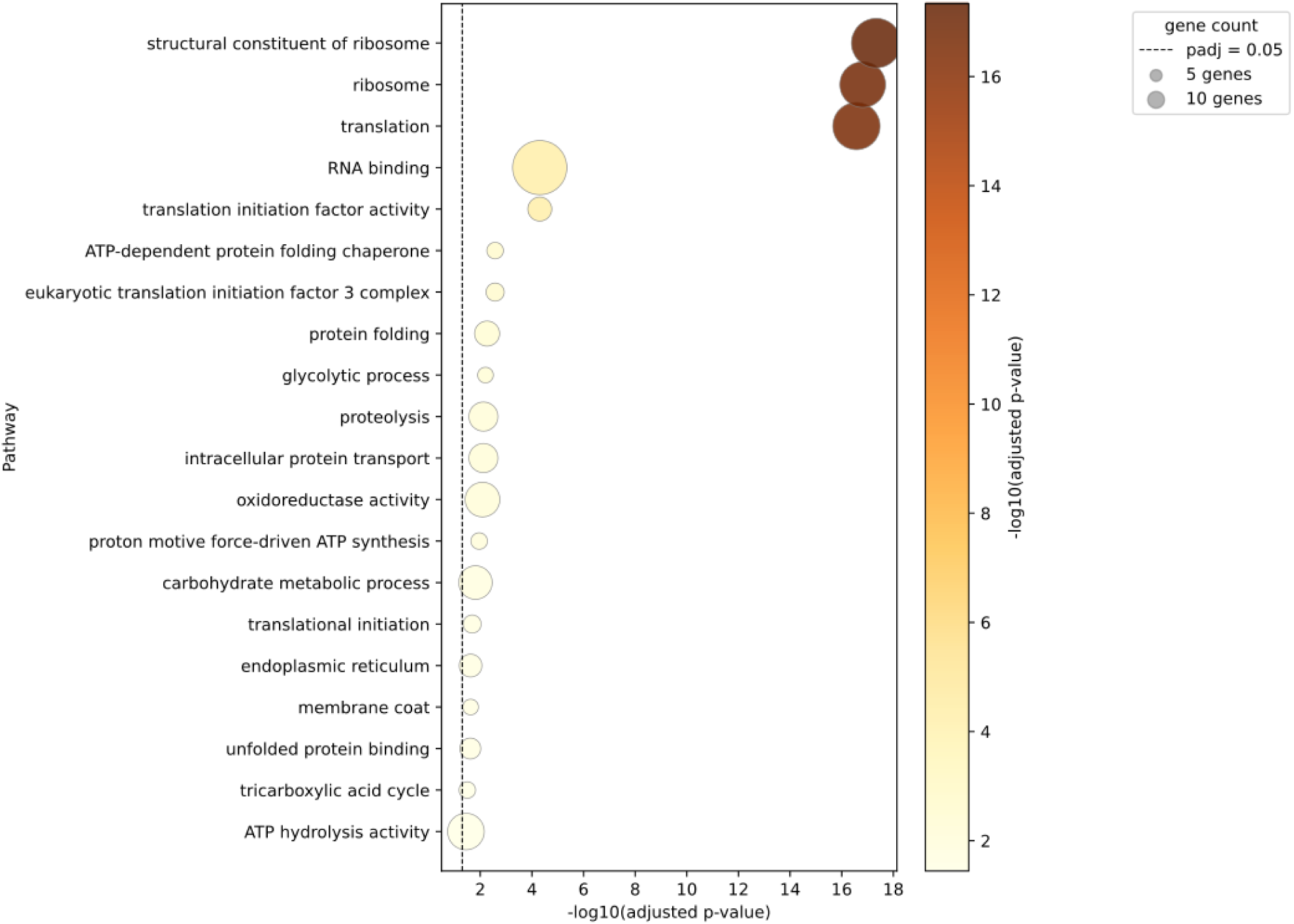
A bubble plot of GO (Gene Ontology) enrichment. The bubble plot summarized the biological functions of genes in the first quadrant of the scatter plot (Fig.14). The x-axis represents the level of enrichment. The y-axis is GO terms sorted by gene counts. Bubble colors represent adjusted p-value. Red means more significant, yellow means less significant. Bubble sizes represent the number of genes clustered by GO terms.

An additional 159 annotated TUM1 genes were included in Iso-Seq-dominated PGs, suggesting that they were grouped with translated protein sequences from the Iso-Seq dataset due to shared peptides, while unique peptides were detected exclusively in their Iso-Seq counterparts. We manually curated the corresponding Iso-Seq-derived predictions and identified 140 annotated TUM1 genes that potentially encode novel proteins from their non-canonical isoforms. We computed proportions of total peptide accessions covered by the canonical genes in these PGs. The histogram was right-skewed and showed a peak between 80% and 100% (excluding 100%) (Fig. 16. These proteins, supported by novel peptides, reflect either improved annotation or novel proteins derived from novel isoforms.

**Figure 16.**
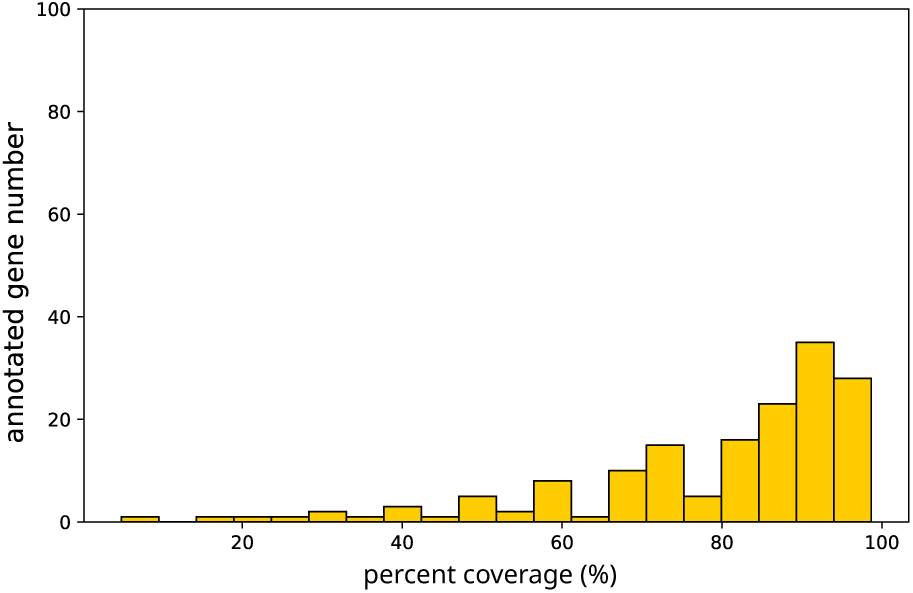
The histogram shows the percentages of peptides covered by annotated TUM1 genes in Iso-Seq-dominated protein groups. Percent coverage is defined as the number of peptides mapped to TUM1 genes relative to the total peptide number (also the peptide number mapped to the leading protein).

In addition to proteomic evidence for canonical TUM1 genes, we also detected 269 predicted proteins from the intergenic space, represented in *Bh* TUM1-related PGs. These 269 proteins originate from TEs (Fig. 17). We found that many of them are nested, but most nested TEs belong to the same TE order; only two are nested across orders. Among these TE-derived proteins, 215 originated from LTRs, 44 from LINEs, 3 from SINEs, and 7 from unclassified retrotransposons or nested retrotransposons across TE orders. Since TEs of the LTR and LINE orders encode proteins required for their proliferation, we further investigated whether the detected proteins correspond to classical TE proteins. To do so, we annotated TE domains for the predicted TE-derived proteins supported by MS using Pfam. We found 68 predicted protein sequences corresponded to classical TE protein domains. 4 predicted TE proteins were near-complete LTR proteins that have Gag protein, integrase, Pol polyprotein, and reverse transcriptase. Other 64 proteins contain incomplete retrotransposon domain arrays. Importantly, within the proteomically supported set of TE proteins, we also detected 201 proteins lacking pre-annotated TE domains, including 3 proteins derived from SINE elements, which are not known to encode proteins. These observations suggest that at least a fraction of proteins originating from intergenic regions may represent novel transposable element–derived proteins.

**Figure 17.**
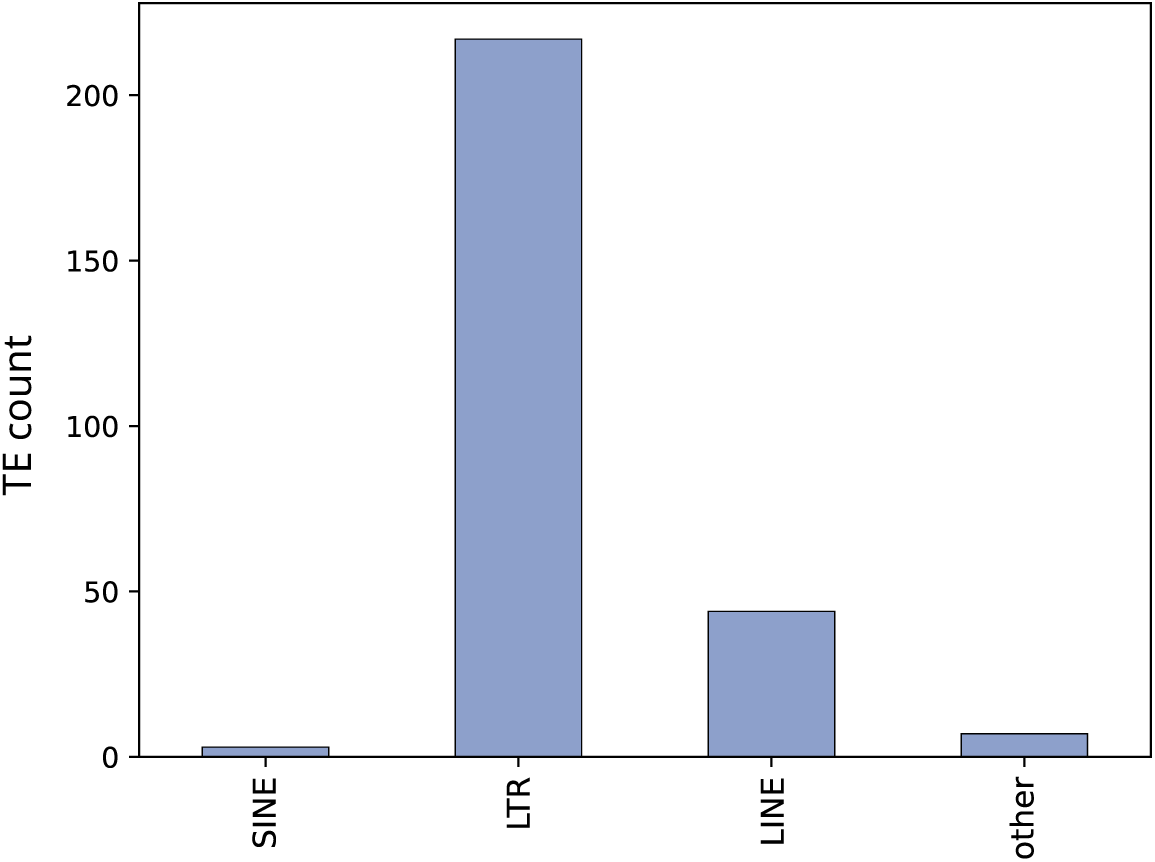
The barplot illustrates the counts of predicted TE proteins identified by MS. The x-axis depicts TE orders whereas y-axis is TE count.

## Discussion

Powdery mildew pathogens have highly complex genomes with a very high transposable element content. As a consequence, only few powdery mildew genomes with resolved chromosomal structure are currently available in the literature, thereby limiting comparative genomic studies in this group of plant pathogens. Here, by combining several long-read sequencing datasets with Hi-C, we were able to resolve chromosomal structures in *B. hordei*. Importantly, the resulting assemblies from both isolates yielded highly similar results, thus supporting the quality of our assemblies. Like *Bh*, both the grapewine powdery mildew fungus (*Erysiphe necator*) and *Bgt*, for which chromosomes have been resolved, possess 11 chromosomes (Müller et al., 2019; Zaccaron et al., 2023). *Bgt* and *Bh* were until recently considered host-specific lineages of the same species (i.e. formae speciales), before being reclassified as distinct species (Liu et al., 2021). Despite otherwise strong gene synteny, we identified several large-scale structural rearrangements between their genomes. Such rearrangements are known to contribute to species divergence by impairing efficient recombination between chromosomes. Therefore, the presence of these large-scale structural rearrangements between *Bh* and *Bgt* supports the classification of the two as separate species. Interestingly, such chromosomal rearrangements have recently also been found between subpopulations within *Bgt* (Sotiropoulos et al., 2025). Future macroevolutionary comparisons may support a better understanding of neutrality or function of such structural variations in host-adaptation or speciation processes.

Apart from an overall genomic conservation, we also observed additional similarities, for instance in overall proportional contribution of the individual TE orders to the genome (Müller et al., 2019). Moreover, as in *Bgt*, the centromeres of *Bh* are occupied by a high proportion of LINE transposons (Müller et al., 2019; Sotiropoulos et al., 2025). In addition, we found that SINEs are located closer to genes than any of the other three TE orders, a pattern that has been observed in previous studies for *Bgt* ((Müller et al., 2019; Sotiropoulos et al., 2025)). Due to the thousands of copies of individual TE families in the genomes of Blumeria pathogens and the availability of only short-read RNA-sequencing data, it has so far not been possible to determine whether close proximity to genes results in higher TE expression. Here, our approach using time-resolved long-read Iso-Seq enabled us to identify thousands of transposable element transcripts and to determine their genomic origin, thereby obtaining a much more fine-grained understanding of TE transcription across the genome. Indeed, we found that TEs that are part of transcripts constituting TE–gene chimeras were expressed at higher levels than their intergenic TE counterparts, providing evidence that positional effects influence TE expression. Among all TE orders, SINEs stood out in particular by containing the highest percentage of expressed TE loci as well as the most intact insertions into UTRs of genes. The reason for the enrichment of SINEs in gene UTRs remains unclear. It is possible that SINEs preferentially insert near genes; alternatively, due to their shorter size, such insertions may be better tolerated, as they are less disruptive. The high frequency of TE-gene chimeric transcripts is further evidence of co-transcriptional activity. Our data now allow for closer inspection of to which extent this is driven by TE read-through or upstream gene promoter activity.

Our deep proteomics approach of a single time point of *Bh* interaction with host epidermal tissue validated a high amount of predicted gene models as protein-coding genes. In total we found peptide evidence for 4,643 genes that showed transcript evidence at 50 hpi. Taking into consideration, that untargeted proteomics has certain technical limitations, and we inspected only one time point in the life cycle of *Bh*, this number appears very high, also compared to earlier proteomics studies in *Bh* (Bindschedler et al., 2009). Data further evidenced peptides from 241 effector candidates, again exceeding previous numbers of 97 identified effector proteins (Pedersen et al., 2012). Sixty-three of those effectors belong to the quadrant that we define as highly expressed both at peptide and transcript quantification levels. Some of them are CSEPs conserved in *Bgt* (Müller et al., 2019), suggesting they are ancient highly expressed effectors with core functions in fungal pathogenic success.

Our combined long-read RNA sequencing and proteomics approach shows that TEs in *Bh* are not only expressed at the transcriptional level but are also translated during the asexual growth phase. While previous reports have provided peptide evidence for the translation of ORF1 from LINE transposons in *Bh* (Amselem et al., 2015), our study reveals translation of hundreds of TE proteins from all three TE orders (LTRs, LINEs, and SINEs). As expected, a significant proportion of the identified TE-derived proteins encode canonical TE proteins involved in TE proliferation. However, we also identified many cases for which no bona fide TE-related protein domains could be predicted. A more detailed and careful analysis will be required to determine whether these proteins represent previously unannotated TE proteins or truly novel proteins derived from TEs. Notably, the detection of proteins encoded by SINE elements, which are not known to possess intrinsic protein-coding potential, indicates that TEs may indeed give rise to novel domesticated proteins. Our study demonstrates that the large TE fraction of the *Bh* genome has the potential to produce novel TE-derived transcripts and proteins. This opens a new possibilities for understanding whether and how individual TE interact with neighboring genes at the level of expression, recombination and other positional polymorphisms (compare also (Sotiropoulos et al., 2025)). Canonial TE transcripts, TE-containing or TE-antisense transcripts might further have gene or TE-regulatory function within *Blumeria* or even in cross-kingdom targeting of host genes (Kusch et al., 2024, 2023; Qian et al., 2023; Kunz et al., 2024). In addition, TE bear a potential for the expression of previously overlooked TE proteins or proteins that newly evolve from TE. These diverse mechanisms together with the high TE content in *Bh* could potentially support adaptation to new host environments that is crucial for an obligate biotrophic parasite such as *Bh*. Closer manual inspections of individual TE-derived innovations in combination with pan-genomics (Sotiropoulos et al., 2025) and a deeper understanding of the selective forces that act on *Blumeria* genomes and TE loci will shed light on potential advantages of carrying a high load of TE in a parasités genome.

## Methods

### Fungal isolates

*Blumeria hordei* isolate TUM1, collected in Germany, has been established for several years at the Chair of Phytopathology, Technical University of Munich, Germany. To maintain a consistent genotype, TUM1 was purified by two rounds of single-spore colony isolation and then propagated on barley leaf segments on plates. TUM1 was cultured and propagated asexually in growth chambers at 20°C and 60% humidity with long-day conditions(16h daylight) on the susceptible barley variety ‘Golden Promise’. The progeny of purified TUM1 was maintained on plates in growth chambers at 4°C with long-day conditions and was propagated every two months.

*B. hordei* isolate AUS1 was isolated in 2023 as a single colony from a population sampled on the susceptible barley variety ‘Stirling’ at the Hermitage Research Station of the Queensland Government’s Department of Primary Industries, Queensland, Australia (S 12°21.79*^′^*, E 6°4.64*^′^*) in April 2017. The isolate was propagated and preserved on 2.5cm segments of young barley leaves (varieties ‘Baudin’ and ‘Stirling’) placed on a benzimidazole agar medium (with 38 mL of 0.5 gr/L benzimidazole/MilliQ water for every 600mL of water) in Petri dishes in an incubator with 12-hour light/12-hour darkness cycles and at 20°C.

### DNA extraction, library preparation and genome sequencing

High-molecular-weight DNA extraction was conducted as described in Bourras et al. (2015). Briefly, conidia spores were collected from propagation plates. 100 mg of spores were flash frozen in liquid nitrogen and ground in a 2ml tube using 3mm stainless-steel beads and a high speed plate shaker. After three rounds of grinding (30 seconds at a frequency of 30/s) 300 µl of a 5% Sarcosyl solution preheated at 65°C were added, followed by vigorous vortexing and immediate supplementation with 700 µl of a solution containing 0.2 M Tris(hydroxymethyl)aminomethane at pH 7.5, 50 mM EDTA, 2 M NaCl, 2% Cetyl trimethylammonium bromide (CTAB) and 0.25 M Sodium metabisulfite (*Na*_2_*S*_2_*O*_5_). For cell disruption, tubes were vortexed vigorously and incubated 15-30 minutes at 65°C. After incubation, 600 µl of Chloroform were added, followed by a 14 000 RCF centrifugation for 10 min at 4°C. The supernatant was carefully collected and supplemented with 1 volume of 100% isopropanol at −20°C. The tubes were mixed by gentle inversion 6-8 times and then centrifuged 10 min at 14 000 rcf, 4°C. After removing the supernatant, the pellet was dried on ice for 15-30 minutes and then re-suspended in 450 µl of standard TrisEDTA buffer. The dissolved pellet was purified using Amicon Ultra 0.5 ml centrifugal filters MWCO 30kDa (Sigma-Aldrich, Steinheim, Germany) according to the manufacturer. The DNA isolation was followed by RNA-ase treatment, by adding 300 µl TE and 4 µl RNAse (DNAse free, 10 mg/ml) and an incubation of 1h at 37°C. The DNA was then re-purified by adding 300 µl Phenol–chloroform and inverting the tubes 6 – 8 times. After 10 min cetrifugation (14 000 RCF at 4°C), the DNA in the supernatant was precipitated by adding 2.5 volume of −20°C 100% Ethanol2 and 0.01 volume 3M AcNa. After incubation overnight at 4°C, the precipated DNA was collected by centrifugation (14 000 RCF at 4°C) and washed with 450 µl 70% Ethanol, followed by a second centrifugation (14 000 RCF at 4°C). The DNA pellet was dried on ice and the suspended in 45 µl TE or water. Whole genome DNA for TUM1 was used for different sequencing technologies: 1) long-read sequencing using Oxford Nanopore Technology (PromethION 24, library preparation with barcoding kit SQK-NBD114-96 following the manufacturers instructions), conducted by the Gene Centre Munich, Germany; 2) short read paired-end sequencing using Illumina NovaSeq™ X Plus (2 x 150 bp) conducted by GENEWIZ Germany GmbH; 3) long read ultra low input PacBio Revio sequencing performed by the Max Planck-Genome-Centre Cologne, Germany (https://mpgc.mpipz.mpg.de/home/). Hi-C libraries for TUM1 were generated from conidia spores using the Arima High Coverage Hi-C kit (Arima Genomics, A410110, Carlsbad, CA, USA) according to manufacturer’s instructions to generate Hi-C paired-end (2 × 150 bp) reads on a Illumina Nextseq 2000 instrument, also performed by the Max Planck-Genome-Centre Cologne, Germany. Genomic DNA of AUS1 was sent to Macrogen in South Korea, where a PacBio HiFi library was created, and the sample was sequenced using half a SMRT cell on the PacBio Revio. The long reads output was used for the subsequent *in silico* analyses. Conidia spores of AUS1 were sent to the Biomolecular Resource Facility (BRF), Australian National University (ANU), Canberra, Australia. Hi-C data was retrieved using the fungal mycelia and the PhaseGenomics Proximo Hi-C (Fungal) kit protocol (KT6040 kit). The Hi-C prepared libraries were used to sequence Illumina paired-end short reads using half a flow cell of NovaSeq X 1.5B (300 cycles run).

### RNA extraction, library preparation and RNA sequencing

Three independent replicates of a time-course infection experiment on barley epidermis were conducted to gain insight into the fungal gene transcription during infection. TUM1 spores were harvested and used to infect the abaxial surface of the first leaf of 8-day old barley seedlings, variety ‘Golden Promise’. The infected epidermis was collected at 1, 10, 20, 30, 40, and 50 hours post-infection (hpi) and was frozen in liquid nitrogen. Each sample was assumed to contain a mixture of polyadenylated RNA that derived from *Bh* and from barley. Total RNA was extracted from the epidermis using the QIAGEN RNeasy Plant Mini Kit, following the manufacturer’s protocol. Genomic DNA contamination was removed using the QIAGEN RNase-Free DNase Set and total RNA was recovered with Amicon Ultra-0.5 (Millipore) centrifugal filters. RNA integrity was assessed by 1% agarose gel electrophoresis, confirming the absence of degradation. RNA concentration was quantified using a Qubit™ fluorometer with the Qubit™ RNA Broad Range (BR) Assay Kit. The RNA sequencing was conducted at Genewiz Germany Gmbh (Leipzig, Germany). All 18 samples (three replicates and six time points) were sequenced using Illumina NovaSeq technology after double rRNA depletion for plant and fungi to enrich mRNA and non-coding RNA. The resulting stranded paired-end reads (2×150 bp) had a depth of 20 million reads per sample. Only four time points (10, 20, 30, 50 hpi) and pooled RNA samples from the replicates for each time point, were sequenced using PacBio Isoform Sequencing (Iso-Seq). For this, PacBio SMRTbell® Iso-Seq HiFi libraries were constructed and sequenced with Sequel IIe system for 24 h movie time. The average sequencing depth reached over 4 million HiFi reads per sample.

### Mass Spectrometry Sample Preparation

The epidermis samples for protein extractions were in the same batch as RNA sequencing experiment. The protocol for plant tissue protein extraction was applied[add appendix] for three independent replicates at 50 hpi, named PA-50, PB-50 and PC-50 and dissolved in 1:10 protein lysate. Cell pellets from three biological replicates (PA_50, PB_50 and PC_50) were pooled and lysed in 4% SDS, 40 mM Tris-HCl (pH 7.6), 1% TFA and 2% NMM. The total protein concentration was determined using the BCA Protein Assay Kit (Thermo Scientific). SP3 magnetic beads (Sera-Mag A and B) were combined in an equimolar ratio, washed three times with deionized water and resuspended again in the starting volume of deionized water. Three technical replicates were performed by three times mixing 50 µl of protein sample (corresponding to 50 µg total protein) with 7.5 µl SP3 bead slurry. Proteins were precipitated by adding EtOH to a final concentration of 70%. Samples were washed thrice with 200 µl 80% EtOH and once with 100% ACN. Reduction and alkylation took place in 100 µl digestion buffer (100 mM HEPES pH 8.5, 10 mM TCEP, 50 mM CAA) for 60 min at 37 °C and 1000 rpm. Trypsin was added in a 1:50 ratio. Samples were incubated overnight at 37 °C and 800 rpm, subsequently acidified with TFA (1% final concentration), desalted via C18 peptide cleanup, pooled, dried and stored at −80 °C. Next, the sample was off-line fractionated using basic reverse phase chromatography. Briefly, the 150 µg peptide digest was reconstituted in 25 mM ammonium bicarbonate (pH 8) and loaded onto a C18 column (XBridge BEH130, 3.5 mm, 2.1 3 150 mm, Waters Corp.). Peptides were eluted with increasing acetonitrile concentration to 96 fractions and subsequently pooled to 32 fractions. All fractions were dried down before LC–MS/MS analysis.

### Genome assemblies

AUS1: We assembled the genome using two methods. Firstly, we assembled the long reads separately with flye (v2.9.3-b1797) (Kolmogorov et al., 2019), using “–pacbio-hifi” and the default for the rest of the parameters. Secondly, after randomly subsampling for 80% of the PacBio HiFi long reads, and then excluding reads shorter than 3kb, using filtlong (v0.2.1) (https://github.com/rrwick/Filtlong), we assembled the remaining reads using Hifiasm (v0.19.8-r603) (Cheng et al., 2021b) and the default parameters. After that, we used quickmerge (v0.3) (Chakraborty et al., 2016), which uses NUCmer (v3.1) (Marçais et al., 2018) to combine these two assemblies and create a better one. Hi-C reads of AUS1 were aligned to the AUS1 assembly using the nf-core/hic pipeline with the –digestion arima option, and scaffolding was performed using YaHS (v1.2.2, (Zhou et al., 2023)) with the following parameters: -q 1 -r 2000,5000,10000,20000,30000,40000 –no-contig-ec.

TUM1: The ONT long-read sequencing data was assembled using the nf-core/genomeassembler pipeline (vs1.1.0, see https://zenodo.org/records/16267903) using profile ‘‘ont_fly’ and parameters ‘genome_size 120 Mb’, ‘short_reads’, ‘trim_short_read’, ‘polish_pilon’, ‘scaffold_longstitch’, ‘scaffold_links’, and ‘lift_annotations’. For the polishing, the Illumina short-read data for TUM1 was provided. The PacBio sequencing reads were randomly subsampled to 900,000 reads and assembled with hifiasm (v0.19.8-r603, (Cheng et al., 2021a)) using the following parameters: −l0 -f0. The hifiasm assembly was scaffolded using the assembly output of the nf-core/genomeassembler with the RagTag scaffold command (Alonge et al., 2022), implemented in the Galaxy Europe server (Community, 2024). Finally, Hi-C reads were aligned to the TUM1 assembly using the nf-core/hic pipeline with the –digestion arima option, and scaffolding was performed using YaHS (v1.2.2, (Zhou et al., 2023)) with the following parameters: -q 1 -r 2000,5000,10000,20000,30000,40000 –no-contig-ec. The chromosome numbering was assigned based on alignment with the *Bgt* genome (Müller et al., 2019) using D-GENIES (Cabanettes and Klopp, 2018a).

We used BUSCO (v6.0.0) (Manni et al., 2021; Tegenfeldt et al., 2025) to check for genome completeness with the ascomycota_odb10 database and ascomycota_odb12 database.

### Comparative analysis

We used whole-genome alignments of TUM1 and AUS1 from NUCmer v3.1 (Marçais et al., 2018) as input for SyRi (Goel et al., 2019) to identify syntenies and structural variations. The rearrangements greater than 100 Kbp were shown on synteny plots with Plotsr (Goel and Schneeberger, 2022).

### SINE curation

The manual curation of SINEs relied on reference genome DH14 (GCA_900239735.1) (Frantzeskakis et al., 2018). 17 *Bgt* and 1 *Bh* SINEs collected from TREP (Wicker et al., 2002)were used as queries and were aligned to DH14 using BLASTn, the package is BLAST+ v2.15.0 (Camacho et al., 2009). The best hits were selected as the homologous SINE representatives in *Bh*. To obtain all family members and their copies in DH14, we searched the genome with BLASTn using *Bh* homologous SINEs as queries. We extracted nucleotide sequences of all hits, including their 100-bp flanking regions, representing variations of SINE copies. Considering the high number of SINEs in the analysis, we did a two-step filtering before the manual process: First, the extracted sequences were clustered by running CD-Hit-Est v4.8.1 to reduce redundant elements (Fu et al., 2012). Second, the remaining 19,858 sequences from 18 query sequences were further filtered using the script ready_for_MSA.sh from Gourbet *et al*., 2022 to extract the 25 most extended sequences and 75 randomly selected sequences for each query (Goubert et al., 2022). Together with the queries, these sequences were passed down to Mafft v7.505 for MSA (Katoh and Standley, 2013). We then visualized the alignments in AliView v1.28 to identify ends of SINEs and remove faulty sequences (Larsson, 2014). We used Trimal v1.4.1 to reduce alignment gaps. TEtrimmer was applied to crop the flanking regions (Qian et al., 2025). SINEs shorter than 200bp, however, were cropped manually. Consensus sequences of SINEs were generated with CONS EMBOSS (Madeira et al., 2024).

### TE classification

TE classification of non-SINE elements followed the strategy from Qian *et al*. by TE domain prediction using Pfam_scan.pl and comparison to the published TE libraries Repbase and TREP (Jurka et al., 2005; Wicker et al., 2002; Qian et al., 2023; Madeira et al., 2024). SINEs were removed before this classification step. The non-SINE TEs were first compared to TREP by BLASTn for classification. The TE was counted as classified and belonging to the same TE family if at least 80% of the length of both query and reference sequences aligned with an identity score greater than 80. TEs that were not found in TREP were similarly compared to Repbase for classification. Following this, a search for tansposase or reverse transcriptase domains, which are considered key domains defining the class of a TE, was conducted for the remaining TEs. TEs that coded for these were grouped as “classified”. If no key domain was predicted, the TE was grouped into “fragmented”. All remaining TEs were grouped into “no domain”. For the “classified” sequences, the classification was manually checked based on the order of domains in TEs according to the definition of TEs (Wicker et al., 2007). “Fragmented” and “no domain” groups were considered unclassified. An attempt was made to identify the superfamily for the sequences in these two groups using the machine-learning software TEClass2, followed by a manual check (Bickmann et al., 2025). The superfamily name was used if the TEClass2 result matched the initial name; otherwise, TEs were simply classified as “Unknown”.

### Iso-Seq data processing

We processed Iso-Seq data of 4 samples from different time points with package isoseq v4.0.0 (https://github. com/PacificBiosciences/IsoSeq). The SMRTbell molecules were processed by CCS v6.4.0 to generate HiFi reads and predicted accuracy is *≥* Q20 (https://github.com/PacificBiosciences/ccs). To get full-length reads, the primers were removed with lima –isoseq –peek-guess (v2.9.0). Poly A tails and concatemers were trimmed with *isoseq refine*. We applied *isoseq cluster*, which performs hierarchical clustering to generate high-quality consensus transcripts and a *cluster_report.csv*, keeping the number of reads clustered in each group. The transcripts were mapped to the concatenated reference genome of *Bh* isolate TUM1 and barley variety Golden Promise using minimap2 v2.17-r941 (Li, 2018). We used the following setting for transcript mapping: *minimap2 -a -k 15 -w 10 –splice -A1 -B6 -z 200 -g 1500 -C 5 -G 3000*. We used SAMtools v1.6 to extract the *Bh* transcripts and re-mapped them to TUM1 (Danecek et al., 2021). The *TAMA collapse* identified transcript models (TMs) by further collapsing aligned transcripts without 5’ cap selection. This is to reduce the effects of 5’ degradation in TM prediction (Kuo et al., 2020). To retrieve the transcript models, we applied *TAMA collapse* with “-x no_cap -e common_ends -d merge_dup” The outputs were used for downstream pipelines to produce ORF/NMD prediction in genome annotation and reference data in proteomics.

### Genome annotation

The genome of TUM1 was masked with the TE library using RepeatMasker v4.1.1 (Smit et al., 2015). Gene models were first predicted using BRAKER3 (Gabriel et al., 2024), MAKER (v2.31.11+galaxy2) (Holt and Yandell, 2011), Liftoff (Shumate and Salzberg, 2021), and TAMA (Kuo et al., 2020). We then established a pipeline to obtain a comprehensive gene prediction based on the four sources. BRAKER3 integrates genomic, RNA-seq, and published fungal protein database evidence. MAKER performed an *ab initio* prediction using the *Bgt* protein database as the protein reference. Liftoff maps gene annotations from the DH14 genome to the TUM1 genome. TAMA generated gene models from Iso-Seq data. To obtain gene prediction from the long-read evidenced transcriptome, we ran *TAMA collapse* on samples from four time points only with a filter for 3’ priming. The outputs were combined using *TAMA Merge* to predict ORF with TAMA. We set parameters as “-e common_ends -d merge_dup”. This configuration produced conservative results and retained as many isoforms as possible from different time points during the merging process. We ran the pipeline for *TAMA GO: ORF and NMD predictions*, which performs a BLASTp (v2.15.0) search against UniProt (release-2024_03) (Consortium, 2019) and predicts ORFs based on the top hits. We then incorporated the results of BRAKER3, MAKER, and Liftoff using InGenAnnot v0.0.11 (Lapalu et al., 2025). TAMA ORF prediction was not included as a prediction but as long-read evidence in InGenAnnot to ensure independent validation. Annotation Edit Distances (AEDs) were computed independently based on short-read RNA sequencing (aed_ev_tr), long-read RNA sequencing (aed_ev_lg), and the Uniprot database (aed_ev_pr). InGenAnnot also selected the best gene models from clusters representing the same gene derived from multiple prediction sources. Additionally, untranslated regions (UTRs) were refined using InGenAnnot based on Iso-Seq evidence. We assess the performance of selection by visualizing the distribution of AEDs based on scatter plots and UpSet plots for gene models selected from different sources. Genes belonging to clusters without Liftoff evidence were specifically examined with scatter plots, which represented potential novel genes absent in the DH14 isolate. Gene models were categorized according to aed_ev_tr and aed_ev_lg values. Following manual inspection in each category, gene models with both aed_ev_tr >= 0.5 and aed_ev_lg >= 0.5 were identified as genes requiring manual curation. Manual curation was performed using the Apollo genome browser (Dunn et al., 2019). To support the annotation of genes not expressed in TUM1, we incorporated additional transcriptomic evidence, including the Nanopore-derived transcriptome of K1 (Qian et al., 2023) and Illumina paired-end RNA-seq data from DH14 at 48 hpi (Frantzeskakis et al., 2018). Gene candidates with ORFs lacking support from legitimate transcripts were discarded. A total of 2,665 unique ORF predictions from long-read RNA sequencing were identified by clustering the TAMA outputs with the manually curated draft annotation using the InGenAnnot clustering algorithm; only 16 of these predictions produced non-transposable element BLASTp hits. The remaining 2,649 predictions were analyzed with SignalP v4.1 (Nielsen, 2017) using thresholds of -u 0.34 and -U 0.34 to detect potential missing effector genes. Additionally, 95 TAMA predictions that overlapped with predictions from other sources on the same strand and covered more than 10% of the query length were manually inspected to remove redundant predictions. The mitochondrial genes were annotated using MFannot (https://megasun.bch.umontreal.ca/apps/mfannot) (Lang et al., 2023).

### Function annotation and effector prediction

We annotated functions of TUM1 genes with InterProscan v5.76 (Jones et al., 2014), KofamScan v1.3.0 (Aramaki et al., 2020), eggNOG-mapper v2.1.13 (Cantalapiedra et al., 2021), SignalP 6.0 (Teufel et al., 2022) and DeepTMHMM 1.0 (Hallgren et al., 2022). Due to the diversity and plasticity of effectors, we performed Markov clustering on HipMCL for all annotated TUM1 protein sequences, candidate secreted effector proteins(CSEPs), and Bgt effectors (Azad et al., 2018; Pedersen et al., 2012; Müller et al., 2019). All-to-all BLASTp was performed with DIAMOND v2.1.16 (Buchfink et al., 2021). Based on a histogram of E-value and sequence coverage of hits, the self hits were removed, and e-value < 1 - e5, coverage < 75% was filtered out, and e-values were used to weigh the edges of genes. A fine-grained clustering was performed with inflation parameter -I 5. We applied EffectorP 3.0 in effector prediction (Sperschneider and Dodds, 2022).

### LC-MS/MS data acquisition

LC-MS/MS data acquisition of all samples was carried out on a Dionex Ultimate 3000 RSLCnano system coupled to an Orbitrap Fusion LUMOS mass spectrometer (ThermoFisher Scientific, Bremen). Injected peptides were delivered to a trap column (ReproSil-pur C18-AQ, 5 µm, Dr. Maisch, 20 mm × 75 µm, self-packed) at a flow rate of 5 µL/min in 0.1% formic acid in HPLC grade water. After 10 minutes of loading, peptides were transferred to an analytical column (ReproSil Gold C18-AQ, 3 µm, Dr. Maisch, 450 mm × 75 µm, self-packed) for one minute, and then separated using a 50 min gradient from 4% to 32% of solvent B (0.1% FA, 5% DMSO in acetonitrile) in solvent A (0.1% FA, 5% DMSO in HPLC grade water) at 300 nL/min flow rate. The Orbitrap Fusion LUMOS mass spectrometer was operated in data-dependent acquisition (DDA) and positive ionization mode. MS1 spectra (360–1300 m/z) were recorded at a resolution of 60,000 using a normalized automatic gain control (AGC) target value of 100% and a maximum injection time (maxIT) of 50 msec. A cycle time of 2 sec was set. Only precursors with charge states 2 to 6 were selected, and dynamic exclusion of 30 sec was enabled. Peptide fragmentation was performed using higher energy collision-induced dissociation (HCD) and a normalized collision energy (NCE) of 30%. The precursor isolation window width was set to 1.3 m/z. MS2 Resolution was 15.000 with a normalized automatic gain control (AGC) target value of 150% and maximum injection time (maxIT) of 22 msec.

### Input database for peptide-spectrum match (PSM)

To generate a reference database for peptide-spectrum matches (PSMs), we combined Iso-Seq data and canonical protein predictions from genome annotations. To conduct sample-specific protein prediction derived from Iso-Seq data, we first filtered the TAMA output of R-50 with a custom Python module, which takes *cluster_report.csv* and the *TAMA collapse* result as input files. Based on the HiFi read numbers, the module counts the read support for TMs and filters out models with total reads *≤* 2. We noticed TAMA can mis-cluster two genes into one gene group. To avoid this pattern, our module also regrouped transcript models. It applies graph-based methods to assign TMs to two gene groups, with any two TMs assigned to the same group if the shorter TM occupies *≤* 70% of the longer TM’s length. To filter the transcript models in a gene group for further PSM, we assign transcript model *T* a weight (*W^T^*) according to

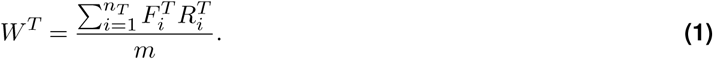

Here, *m* is the total number of unique reads in a gene group. *n* is the number of *T* in a gene group. *R^T^* is the number of reads supporting a transcript. *F^T^* is a fraction assigned to a transcript according to

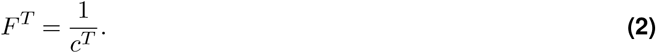

Here, *c^T^*is the number of transcript models the same transcript supports. From the module, we obtained the representative mildew transcript coordinates. We implemented custom scripts to extract transcript sequences from exons and to predict proteins by three-frame translation, filtering for sequences at least 25 amino acids long. The Python modules also computed the genomic coordinates of the predicted proteins. In a previous test with DH14 as the reference genome, we found that the expressed proteins in the Iso-Seq dataset are not significantly correlated with the expressed proteins retrieved from the proteomic dataset. So the input database for MaxQuant is a combination of Iso-Seq and genome annotation. The canonical protein sequences were blasted against the predicted proteins from Iso-Seq data using BLASTp as quality control. The Iso-Seq-derived protein predictions were concatenated with the canonical protein prediction of reference genomes TUM1 and Golden Promise and used as the reference database for PSM.

### LC-MS/MS data analysis

Peptide identification and quantification were performed using the software MaxQuant (version 1.6.3.4) with its built-in search engine Andromeda. MS2 spectra were searched against the fasta files, supplemented with common contaminants, using the built-in option in MaxQuant (Tyanova et al., 2016). The search was performed with two fasta files, including Golden_Promise.protein together with proteome_for_MS_v2. Here the Golden_Promise.protein is the coding sequences of annotated barley (’Golden Promise’ cultivar) genes (Jayakodi et al., 2024). The FASTA file proteome_for_MS_v2 is the hybrid database of TUM1 gene annotation and predicted coding sequences from Iso-Seq dataset. Trypsin/P was specified as a proteolytic enzyme. Carbamidomethylated cysteine was set as a fixed modification. Oxidation of methionine and acetylation at the protein N-terminus were specified as variable modifications. Results were retained at a 100% false discovery rate for Prosit rescoring (Gessulat et al., 2019). The minimal peptide length was defined as 7 amino acids, and the “match-between-runs” functionality was enabled (matching time window 0.7 min, alignment time window 20 min). Mass spectrometry data were further processed using a workflow integrating peptide rescoring, FDR control, and protein-level quantification. Required inputs included evidence.txt, msms.txt, and the protein database FASTA files. Raw files were converted to mzML using ThermoRawFileParser, and peptides were rescored with software Oktoberfest using deep learning-based Prosit_2020_intensity_HCD intensity and Prosit_2019_irt retention time models (Picciani et al., 2024). Rescored PSMs were merged with the original MaxQuant evidence, followed by 1% peptide-level FDR filtering. Protein inference and quantification were performed using the picked group, with 1% protein-level FDR applied.

### Data processing based on Prosit prediction

Quality control and classification of peptide table and protein groups were conducted with custom Python (v3.13) scripts.

### Synteny analysis between *Bgt* and *Bh*

Whole-genome alignment between the *Bh* TUM1 assembly and the *Bgt* CHE_96224 assembly (Müller et al., 2019) was performed using the D-Genies web server with Minimap2 alignments run under default parameters (Cabanettes and Klopp, 2018b). Single-copy orthologous genes among TUM1, AUS1, and CHE_96224 were identified using OrthoFinder (v3.0.1b1) (Emms and Kelly, 2019). Syntenic relationships among these genes were subsequently visualized using the R package circlize (https://github.com/jokergoo/circlize).

## Data availability

All raw data generated in this study are deposited in publicly accessible repositories as follows: Genome and transcriptome sequencing datasets for isolate TUM1 are available through the European Nucleotide Archive (ENA) under accession number PRJEB110634. Genomic sequencing data for isolate AUS1 are deposited in the NCBI Sequence Read Archive (SRA) under accession number PRJNA1443743. The TUM1 and AUS1 assemblies and the corresponding annotation are available at https://gitlab.plantmicrobe.de/marimuel/bh_genome_assemblies_2026.

## Funding

This project was funded by the German Research Foundation (DFG) - TRR 356/1 2023 – 491090170 subproject A03 to Ralph Hückelhoven and Aurélien Tellier. Marion C. Müller acknowledges the support by the German Research Foundation (DFG) grant 543878450.

## Acknowledgment

We would like to acknowledge the technical assistance of Africa Fabro Espada from the Chair of Phytopathology, TUM in propagating *Bh* isolats from the Chair of Phytopathology, TUM. We further acknowledge project I01 of the TRR PlantMicrobe for assistance with data management, including the implementation of FAIR data principles and the facilitation of data and code sharing. We are also grateful to Stefan Kusch and Jiangzhao Qian for valuable discussions on transposable elements as well as Amit Fenn, Luisa Teasdale and Qussai Abbas for discussion on analysis of Iso-Seq and proteomics data. We would like to thank Niklas Shandry for help in implementing the nf-core/genomeassembler pipeline.

## Supplementary Figures

**Figure S1.**
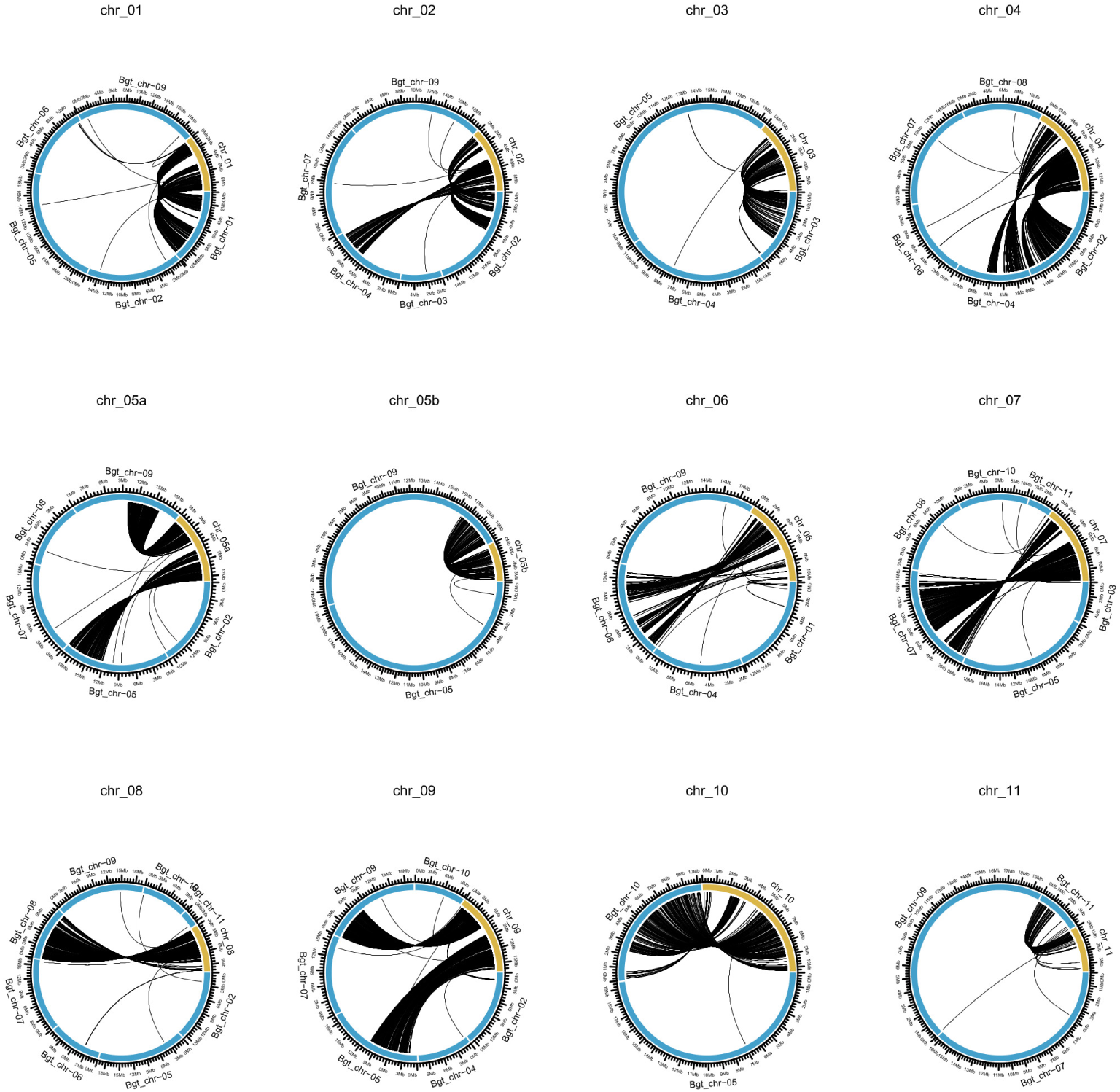
Synteny analysis between Bh isolate TUM1 and Bgt isolates CHE_96224 based on genes conserved in both species. Circos plots for each of the chromosome or chromosome level scaffolds in the TUM1 genome assembly are shown. Position of conserved single copy genes on the chromosome of TUM1 (indicated in yellow) and the corresponding position on the Bgt chromosomes (indicated in blue) are shown.

**Figure S2.**
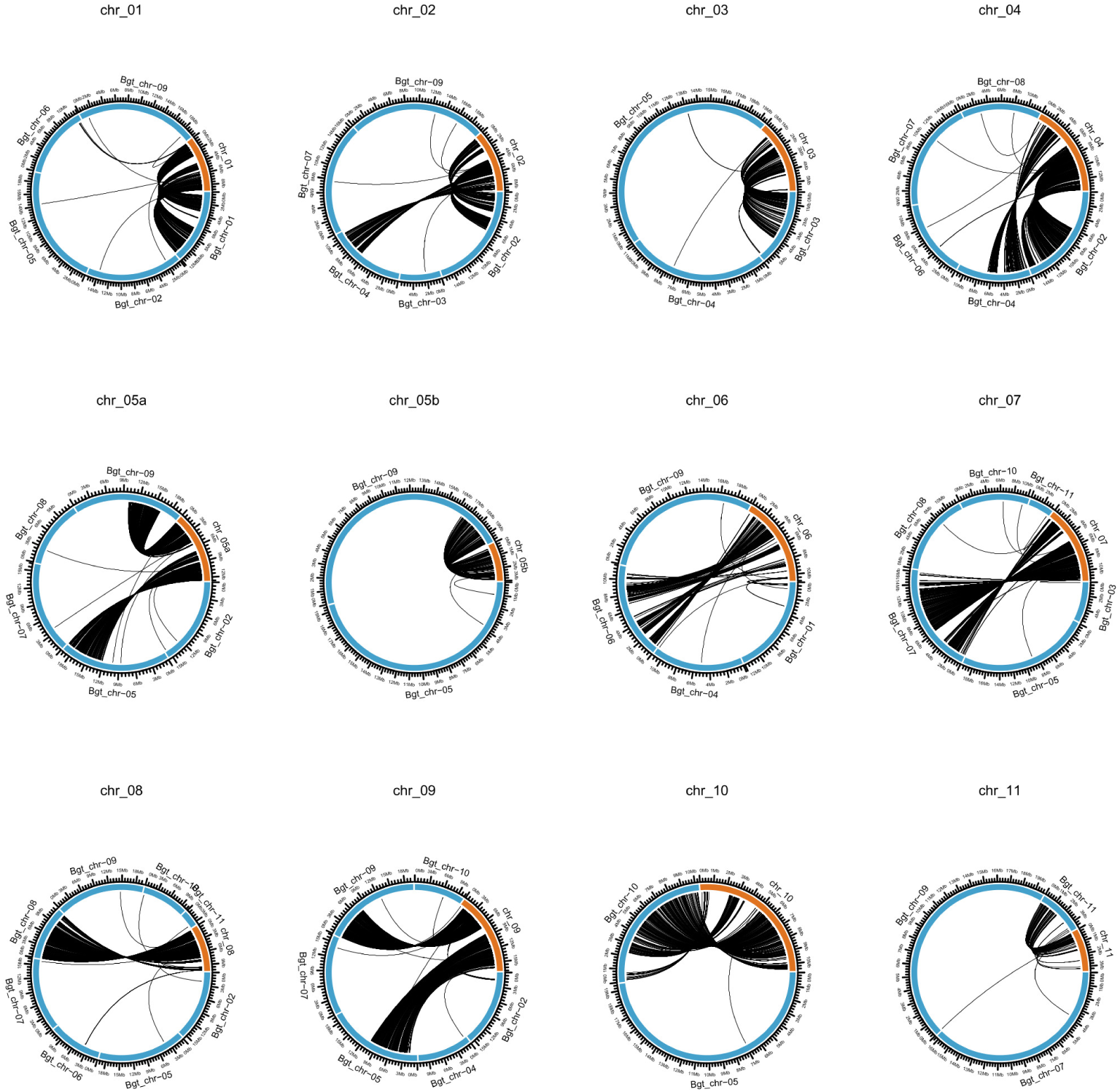
Synteny analysis between Bh isolate AUS1 and Bgt isolates CHE_96224 based on genes conserved in both species. Circos plots for each of the chromosome or chromosome level scaffolds in the AUS1 genome assembly are shown. Position of conserved single copy genes on the chromosome of AUS1 (indicated in orange) and the corresponding position on the Bgt chromosomes (indicated in blue) are shown.

**Figure S3.**
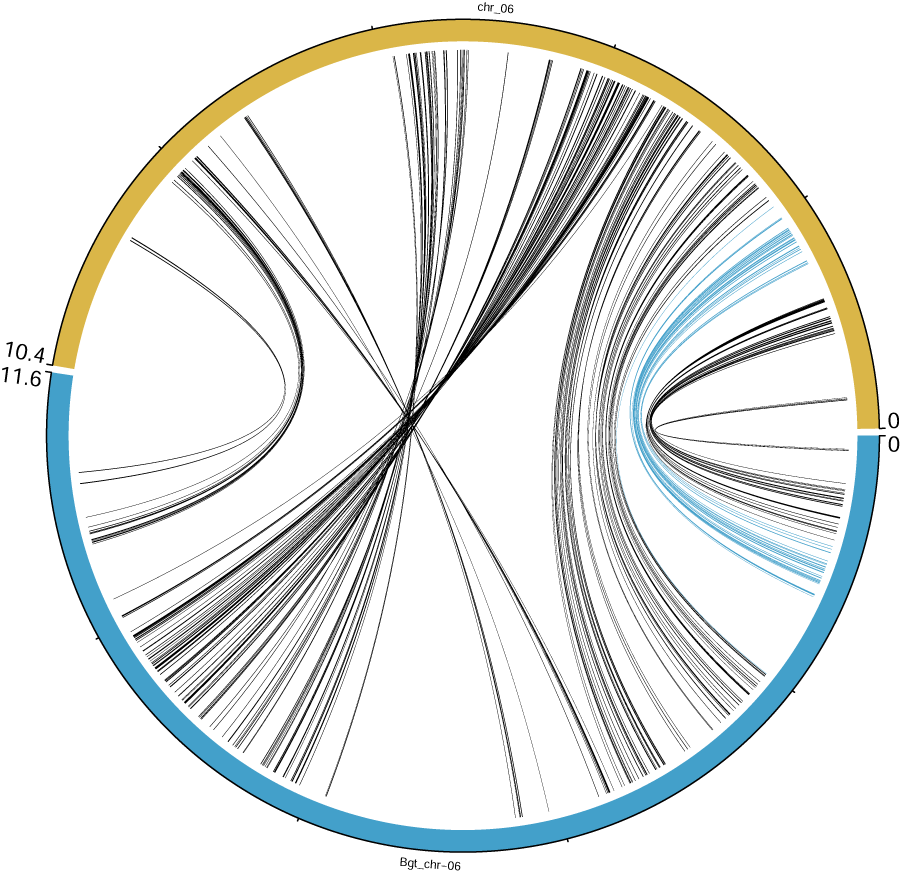
Synteny analysis between Chr-06 of Bh isolate TUM1 (in yellow) and Bgt isolates CHE_96224 (in blue) based on genes conserved in both species. Position of conserved single copy genes on both chromosome are indicated by connecting lines. Blue lines indicates 28 gene located on the inversion that was identified between Bh AUS1 and TUM1

**Figure S4.**
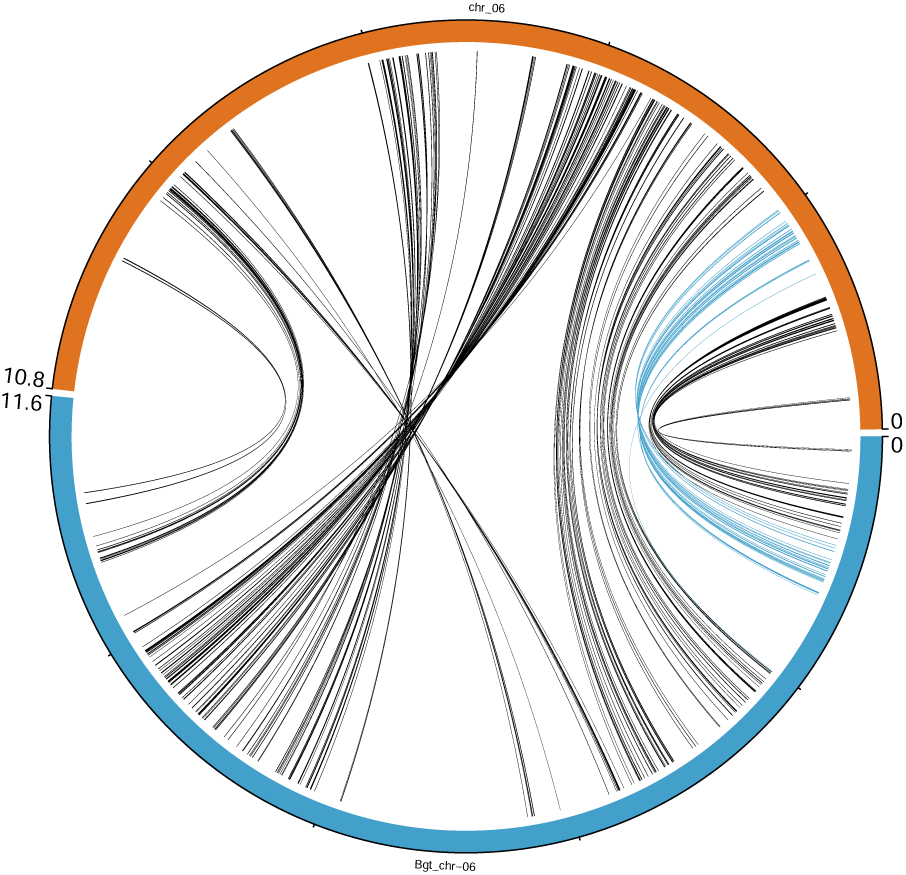
Synteny analysis between Chr-06 of Bh isolate AUS1 (in orange) and Bgt isolates CHE_96224 (in blue) based on genes conserved in both species. Position of conserved single copy genes on both chromosome are indicated by connecting lines. Blue lines indicates 28 gene located on the inversion that was identified between Bh AUS1 and TUM1

**Figure S5.**
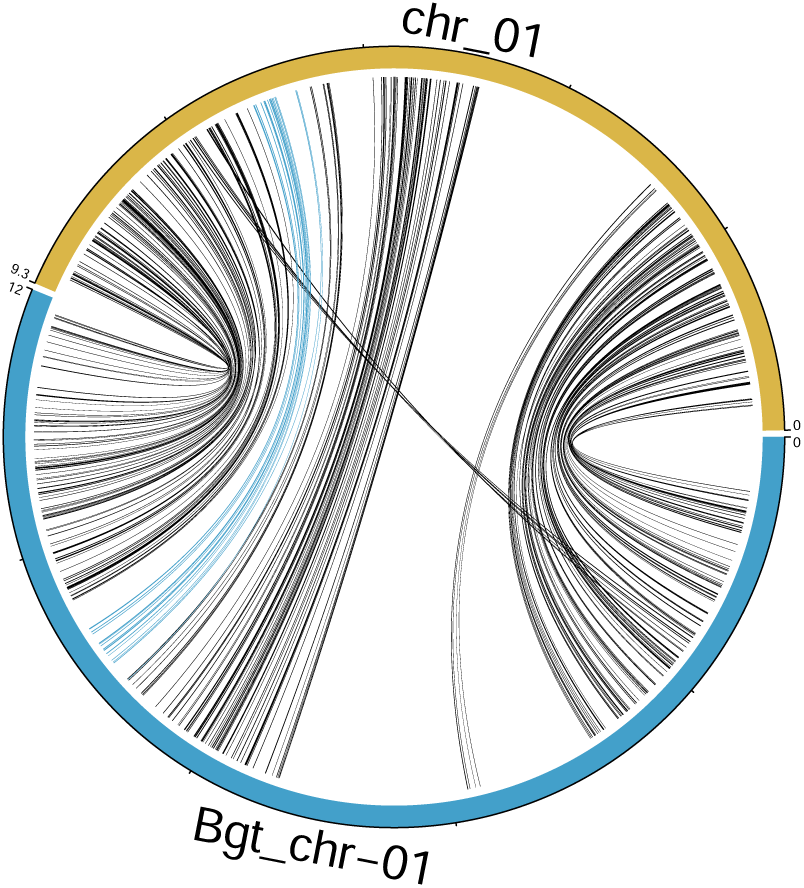
Synteny analysis between Chr-01 of Bh isolate TUM1 (in yellow) and Bgt isolates CHE_96224 (in blue) based on genes conserved in both species. Position of conserved single copy genes on both chromosome are indicated by connecting lines. Blue lines indicates 22 gene located on the inversion that was identified between Bh AUS1 and TUM1

**Figure S6.**
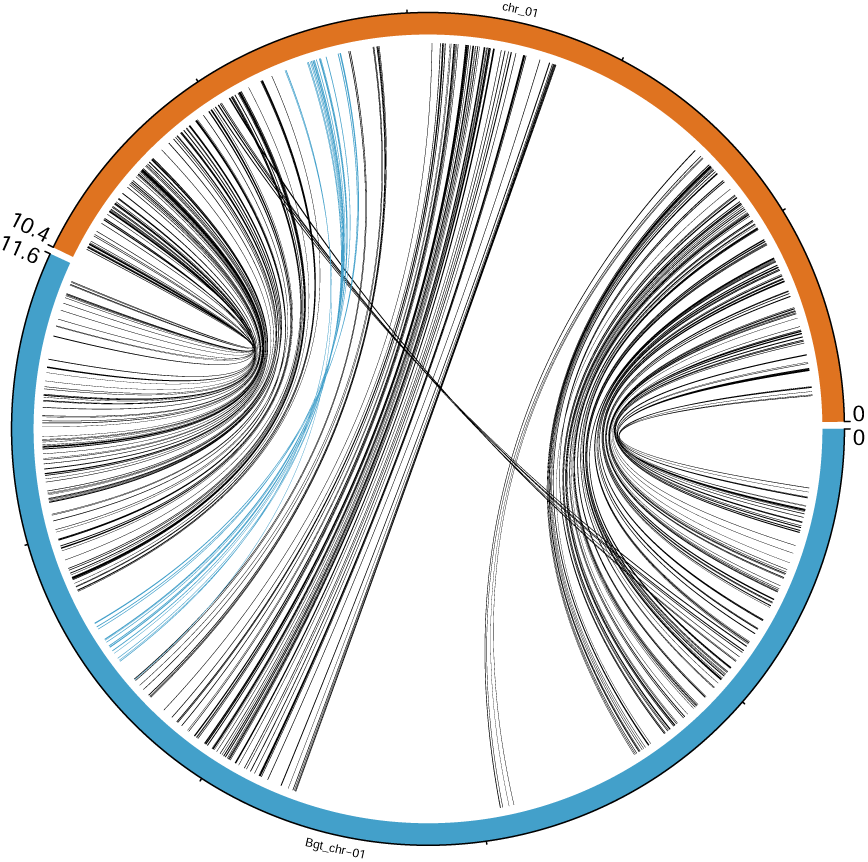
Synteny analysis between Chr-01 of Bh isolate AUS1 (in orange) and Bgt isolates CHE_96224 (in blue) based on genes conserved in both species. Position of conserved single copy genes on both chromosome are indicated by connecting lines. Blue lines indicates 22 gene located on the inversion that was identified between Bh AUS1 and TUM1

## Supplementary Tables

**Table S1.** The mean proportion of total short-read RNA reads mapping to transposable elements (TEs).

| Order | 10 hpi | 20 hpi | 30 hpi | 50 hpi |
| --- | --- | --- | --- | --- |
| LINE | 1.28% | 1.43% | 1.31% | 1.59% |
| LTR | 2.23% | 2.29% | 2.08% | 2.21% |
| SINE | 1.67% | 2.00% | 1.85% | 1.78% |
| TIR | 0.01% | 0.02% | 0.01% | 0.02% |
| sum | 5.19% | 5.74% | 5.25% | 5.60% |

**Table S2.** Summary of structural variation in TUM1 and AUS1.

| Variation type | Count | Length TUM1 (bp) | Length AUS1 (bp) | Proportion (TUM1) | Proportion (AUS1) |
| --- | --- | --- | --- | --- | --- |
| Syntenic regions | 4155 | 97,222,191 | 96,936,899 | 81.97% | 83.01% |
| Inversions | 173 | 4,926,727 | 4,885,077 | 4.15% | 4.18% |
| Translocations | 1778 | 6,102,277 | 6,006,035 | 5.14% | 5.14% |
| Duplications (reference) | 4409 | 10,218,761 | — | 8.62% | — |
| Duplications (query) | 6254 | — | 9,188,747 | — | 7.87% |
| Not aligned (reference) | 2642 | 1,898,240 | — | 1.60% | — |
| Not aligned (query) | 2980 | — | 672,234 | — | 0.58% |

## Notes

### Competing Interest Statement

The authors have declared no competing interest.

